# The Equilibrium Theory of Biodiversity Dynamics: a general framework for scaling species richness and community abundance along environmental gradients

**DOI:** 10.1101/2023.07.17.549376

**Authors:** Jordan G. Okie, David Storch

**Affiliations:** School of Earth and Space Exploration, Arizona State University, Tempe, USA; Center for Theoretical Study, Charles University and the Czech Academy of Science, Prague, Czech Republic; Department of Ecology, Faculty of Science, Charles University, Prague, Czech Republic

**Keywords:** speciation, extinction, ecological limits, latitudinal diversity gradient, community size, biodiversity-ecosystem function relationship

## Abstract

Large-scale temporal and spatial biodiversity patterns have traditionally been explained by multitudinous particular factors and a few theories. However, these theories lack sufficient generality and do not address fundamental interrelationships and coupled dynamics between resource availability, community abundance, and species richness. We propose the Equilibrium Theory of Biodiversity Dynamics (ETBD) to address these linkages. According to the theory, equilibrium levels of species richness and community abundance emerge at large spatial scales due to the population size-dependence of speciation and/or extinction rates, modulated by resource availability and the species abundance distribution. In contrast to other theories, ETBD includes the effect of biodiversity on community abundance and thus addresses phenomena such as niche complementarity, facilitation, and ecosystem engineering. It reveals how alternative stable states in both diversity and community abundance emerge from these nonlinear biodiversity effects. The theory predicts how the strength of these effects alters scaling relationships between species richness, (meta)community abundance, and resource availability along different environmental gradients. Using data on global-scale variation in tree species richness, we show how the general framework is useful for clarifying the role of speciation, extinction and resource availability in driving macroecological patterns in biodiversity and community abundance, such as the latitudinal diversity gradient.

## Introduction

The biodiversity, abundance and biomass of living systems vary extraordinarily across the biosphere, through Earth history, and among major phyla. Understanding this large-scale variation and its consequences for the functioning and resilience of ecological systems is crucial for advancing ecological theory and forecasting the future of biodiversity and ecosystems in the Anthropocene (McGill 2019; Storch et al. 2022). Given the elevated extinctions underway in the Anthropocene, identifying the causes of global-scale patterns of species richness, community abundance and biomass is of particular importance.

The underlying causes for even the most well-studied patterns still remain hotly contested (Pontarp et al. 2019). The pattern of decrease in the number of species from the equator to the poles, documented in nearly all major taxa of multicellular eukaryotes, is one prominent example. Dozens of hypotheses have been formulated (Fine 2015), but no clear consensus has emerged. This lack of consensus is in part due to the multitudinous factors at play but also due to the lack of general theoretical frameworks to test hypotheses and disentangle large-scale diversity drivers.

Recently, evidence is accumulating that large-scale spatial diversity patterns tend to converge onto similar relationships with resource availability and climatic variables regardless of differences in diversification histories (Field et al. 2009; Hawkins et al. 2012; Rabosky 2022), and the origination and extinction rates underlying diversity dynamics exhibit a dependence on diversity (Rabosky 2009; Rineau et al. 2022). These findings have been interpreted as evidence of the role of region or biome-specific ecological limits to species richness (Rabosky and Hurlbert 2015; Etienne et al. 2019), but the nature of these limits has not been entirely clear.

We’ve argued that such limits should be understood as stable equilibria of diversity dynamics that emerge from the links between macroevolutionary dynamics of speciation and extinction modulated by resource availability and community assembly, and that such equilibria shape large-scale spatial and temporal patterns of diversity (Storch and Okie 2019). General theories of biodiversity dynamics (MacArthur and Wilson 1967; Hubbell 2001; Worm and Tittensor 2018) predict the existence of such biodiversity equilibria but suffer from some key shortcomings preventing them from providing a general framework to understanding diversity dynamics. First, they make overly stringent and unrealistic assumptions about the involved processes. For example, the neutral theory of biodiversity, besides assuming ecological equivalence among species that includes equal access to all resources, implicitly assumes that per-species speciation rate increases linearly with population size (Allen and Savage 2007) despite theoretical evidence that speciation rate can exhibit more complex dependencies on species abundance (Gavrilets 2003).

Second, most crucially, while these theories often address (implicitly or explicitly) how community abundance (total biomass or number of individuals) limits species richness by influencing rates of extinction and origination (e.g., Hubbell 2001; Allen and Savage 2007), they take community abundance as an external parameter or static variable and do not address the possibility that the size of the community may itself be a dynamical variable that depends both on the environment and the realized species richness of the community. This is highly problematic since the number of species in a community tends to positively affect the total amount of resources used and converted into community abundance (Loreau 1998; Nijs and Roy 2000; Tilman et al. 2001; Cardinale et al. 2006), as shown in experimental studies of the biodiversity-ecosystem function relationship (a.k.a., the BEF relationship; Bell et al. 2005; Cardinale et al. 2006; O’Connor et al. 2017), suggested by macroecological and paleobiological analyses (e.g., Grace et al. 2016; Liang et al. 2016), and demonstrated with theoretical models (Loreau 1998; Liang et al. 2015). We hereby refer to this effect as the biodiversity effect on community abundance (BECA).

We contend that there is a need for theory that addresses the coupled dynamics of species richness and community abundance. Such coupled dynamics may have non-trivial consequences for biodiversity patterns. The bidirectional interactions between biodiversity and community abundance could lead to an emergence of multiple equilibria and alter how equilibria respond to changing conditions and environmental gradients. These effects could shape temporal and geographic patterns of biodiversity and biomass, as well as their response to environmental perturbations.

Here we provide the first formal presentation of the Equilibrium Theory of Biodiversity Dynamics (ETBD) and present a new quantitative framework for understanding changes in species richness and community abundance along environmental gradients at large spatial and temporal scales. We build on and substantially add to previous work in which we first described the bones of the theory (Storch et al. 2018, 2022; Storch and Okie 2019). The new framework predicts the scaling relationships between equilibria of biodiversity and community abundance and their variation along gradients in extinction, speciation, and resource availability, addressing (but not assuming) the possibility that species diversity may influence resource consumption and community abundance.

The theory also addresses the role of the species abundance distribution and of the population size-dependence of extinction and speciation in modulating the equilibria. It demonstrates how alternative equilibria in biodiversity and community abundance naturally emerge from few assumptions. To illustrate the framework’s application, we apply it to global-scale variation in tree diversity, shedding light on the underpinnings of the latitudinal diversity gradient in trees.

## Results

ETBD focuses on the dynamics of biodiversity at the scales at which variation in speciation rate outweighs colonization rate in its contribution to biodiversity dynamics. It thus addresses the species richness of large-scale eco-evolutionary units, such as biomes, evolutionary arenas (sensu Jetz and Fine 2012), and biogeographic provinces (sensu Rosenzweig 1995). It considers the species richness and community abundance of biomes to be independently determined by the drivers of biodiversity dynamics such that biomes are assumed to not be samples of each other. Here we use the word *community* to refer to the assemblage of species in these large-scale areas.

### Universal Emergence of Diversity Equilibria

The central assumption of the theory is that a population’s probability of extinction and/or speciation exhibit some level of dependence on population size. There is a solid empirical and theoretical basis for this assumption, in particular that extinction probability is generally negatively dependent on population size (e.g., Pimm et al. 1988; McDonald and Brown 1992; Rosenzweig 1995; Griffen and Drake 2008; Okie and Brown 2009; Ovaskainen and Meerson 2010). A consequence of this assumption is that for communities that aren’t on their way to extinction, the curve quantifying the population-size (*N*) dependence of extinction must necessarily intersect the curve quantifying the N-dependence of speciation. The reason is that the speciation rate cannot be higher than the extinction rate over all values of *N*, since as *N* approaches 0, extinction probability necessarily approaches 1 whereas speciation probability approaches zero. Consequently, at this universal intersection, the (negative) slope of the extinction curves must always be lower than the slope of the speciation curve (Figure 1A).

**Figure 1.**
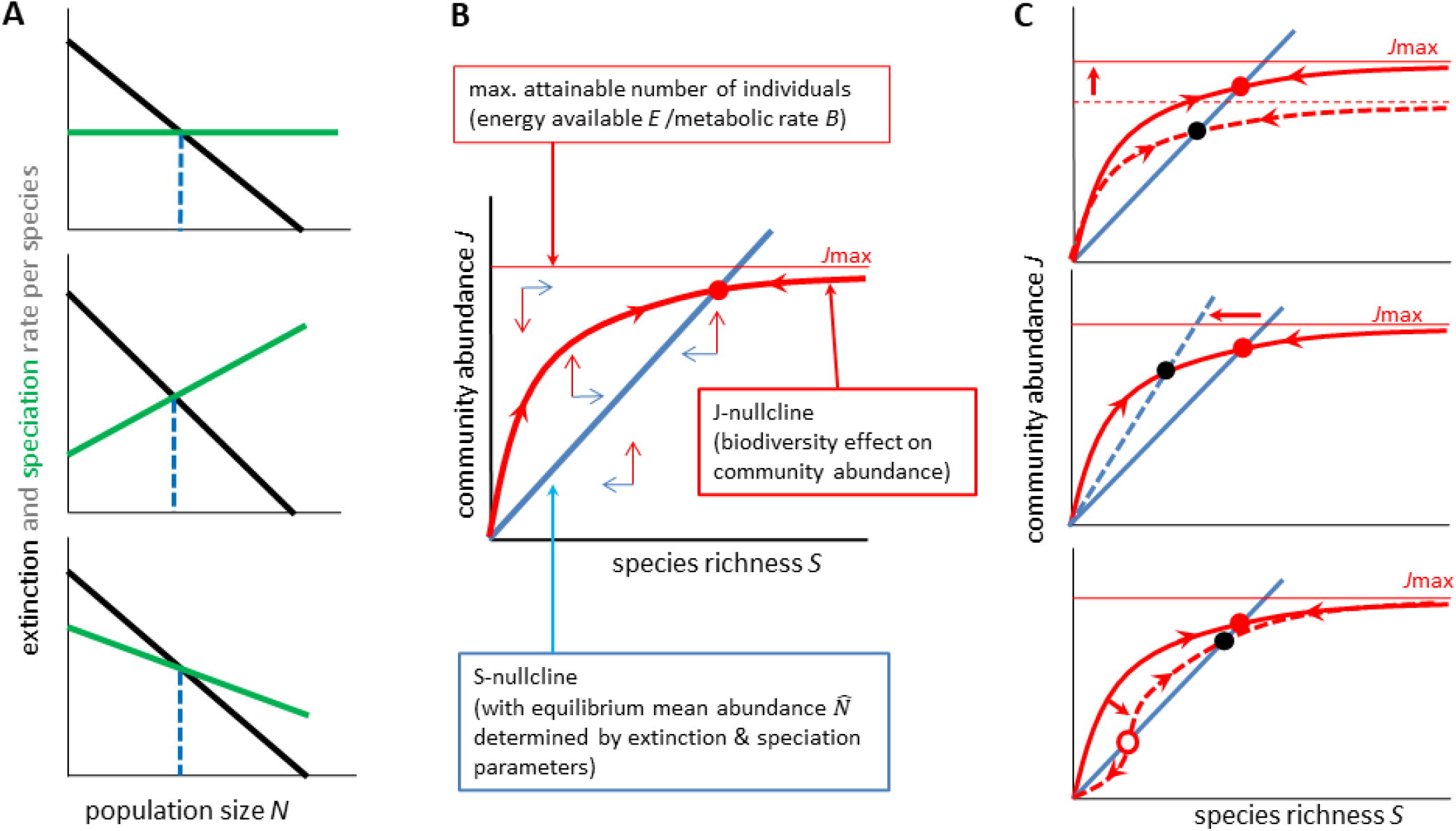
The foundations of the theory. (**A**) Speciation and extinction rates depend on population size in various ways, but around their intersection point the extinction curve (in green) has a more negative slope than the speciation curve (in black). This intersection indicates a population size at which speciation and extinction rates are equal. (**B**) In the phase plane for species richness *S* and by equilibrium mean population size 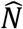, which depends on the intersection of the lines in panel A and the species abundance distribution and indicates values of *S* and *J* in which total speciation rate is equal to total extinction rate. Points to the right of this line represent communities where 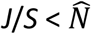; mean population size is thus below 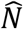, leading to extinction rate being higher than speciation rate and the system having the tendency to move back towards the S-nullcline. The opposite happens if 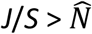 (left from the *S*-nullcline). The equilibrium point for *S* and *J* occurs at the intersection (solid red point) of the S-nullcline with the *J*-nullcline (thick red line), which quantifies the effects of biodiversity on community abundance (BECA). The J-nullcline eventually plateaus, being constrained by energy availability (thin red line). Out-of-equilibrium communities below the J-nullcline follow upward trajectories converging on the J-nullcline, whereas communities above the J-nullcline follow downward converging trajectories. Once on the J-nullcline, trajectories flow along the J-nullcline in the direction determined by their position with respect to the S-nullcline. (**C**) The position of equilibria can change with changing (1) energy availability (top), (2) speciation and/or extinction rates that affect 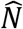 and thus the S-nullcline (middle), or (3) the shape of the J-nullcline (bottom). If the J-nullcline has a more complex shape, there can be multiple equilibria, both stable and unstable (empty circle).

This intersection of curves indicates a population size at which per species speciation and extinction rates are equal. This intersection, along with the form of the probability distribution characterizing the species abundance distribution (e.g., a lognormal distribution or logseries distribution), thus determines the mean species abundance at which a community’s total speciation rate is equal to total extinction rate, which we call the equilibrium mean abundance 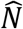. If all species in the community had the same abundance, then 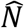 would be exactly equal to the intersection of the speciation and extinction curves. If, instead, species vary in their abundance, then 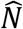 deviates from the intersection, but we have found that it is still set by the intersection and remains uniquely determined by the species-abundance distribution—specifically, the form of the probability distribution of species abundances (see below and SOM). Consequently, in a plot of species richness *S* against community abundance *J* on the vertical axis, there is a straight diagonal line called the S-nullcline that has a slope equal to mean equilibrium population size 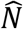 (since the slope is equal to *J*/*S*, which is the mean population size) and represents communities in which total speciation rate is equal to total extinction rate (i.e., dS/dt = 0; Fig 1B, see Supplementary Materials for mathematical details). The equation for this S-nullcline is 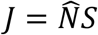. A linear S-nullcline reflects the baseline scenario of biodiversity dynamics in which the parameters of extinction probability, speciation probability, and the probability distribution of species abundances are not directly affected by diversity or community abundance.

Because the negative slope of the extinction curve is lower than the slope of the speciation curve, this nullcline exhibits *attractor* dynamics in which communities perturbed out of equilibrium return back to the S-nullcline, which we thus call an *attractor* S-nullcline. The reason is that points to the right of this S-nullcline represent communities where mean population size (i.e., *J*/*S*) is lower than 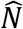 (and thus extinction is higher than speciation rate), whereas in communities located to the left of the S-nullcline, mean population size is higher than 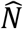(and thus extinction is lower than speciation rate).

### Addressing the Biodiversity Effect on Community Abundance

All points along the S-nullcline represent potential biodiversity equilibria and so a community’s equilibrium point along its S-nullcline is determined by its community abundance *J*. In contrast to other theories of biodiversity dynamics that typically assume that *J* is constant for a community or metacommunity (e.g. Hubbell 2001), ETBD assumes that *J* may itself be influenced by *S*. The rationale for this assumption lies in the realization that *J* is not a feature directly determined by the environment, but instead it is driven by the interplay of population dynamics of all involved species. While in some specific cases *J* may be independent of *S*, it is reasonable to assume an addition of species often leads to the utilization of previously unused resources, and thus S has a positive effect on *J*. Consequently, the J-nullcline, the line delineating the values of *S* and *J* for which *J* can be in equilibrium (i.e., for which d*J*/d*t* = 0), is an increasing function of *S* (Fig 1B thick red line), up to a maximum *J* determined by the maximum energy available to the system divided by mean individual metabolic rate (thin red line, see figure 1B). A community thus has an equilibrium point for *S* and *J* ([*Ŝ, Ĵ*]) located at the intersection of the S-nullcline and J-nullcline.

Whereas theoretical and empirical understanding of the overall shape of the J-nullcline is limited, there are general quantitative features of the J-nullcline that have implications for biodiversity equilibria. All-else-being equal, greater total energy availability increases the total number of individuals that a community of a given *S* can support (figure 1C, top). Consider also that the shape of the J-nullcline reflects the balance of ecological factors governing the slope of the J-nullcline at a given *S* and the degree to which these factors change or not with *S*. At a given *S*, the slope *β* of the line tangent to the J-nullcline in log-log space—that is, *β* = *d*[*log*(*J*)]/*d*[*log*(*S*)]— quantifies the aggregate effects of increasing species richness on the consumption and conversion of resources into abundance (figure 2). We consider three general, fundamental scenarios. (1) Facilitation and ecosystem engineering (*β* > 1): When positive effects of diversity on resource use dominate, each addition of a species increases, on average, the mean abundance at which dJ/dt = 0, leading to an upward-acceleration in the J-nullcline. (2) Ideal complementarity (*β* = 1): When each additional species is, on average, able to use a whole new set of resources just as effectively as the present species and without taking resources away from the present species, the J-nullcline follows a straight line. (3) Competition with some level of niche complementarity (0 < *β* < 1) or no niche complementarity, i.e. complete niche overlap (*β* = 0): If each additional species converts fewer unused resources into abundance than the present species, then the J-nullcline exhibits decelerating behavior. This behavior may have several reasons, including niche overlap or a need to specialize due to increasing competition, and is expected when the community approaches energetic limits of the environment, i.e. the upper ceiling to community abundance. In general, values of *β* close to zero indicate more intense competition for resources.

**Figure 2.**
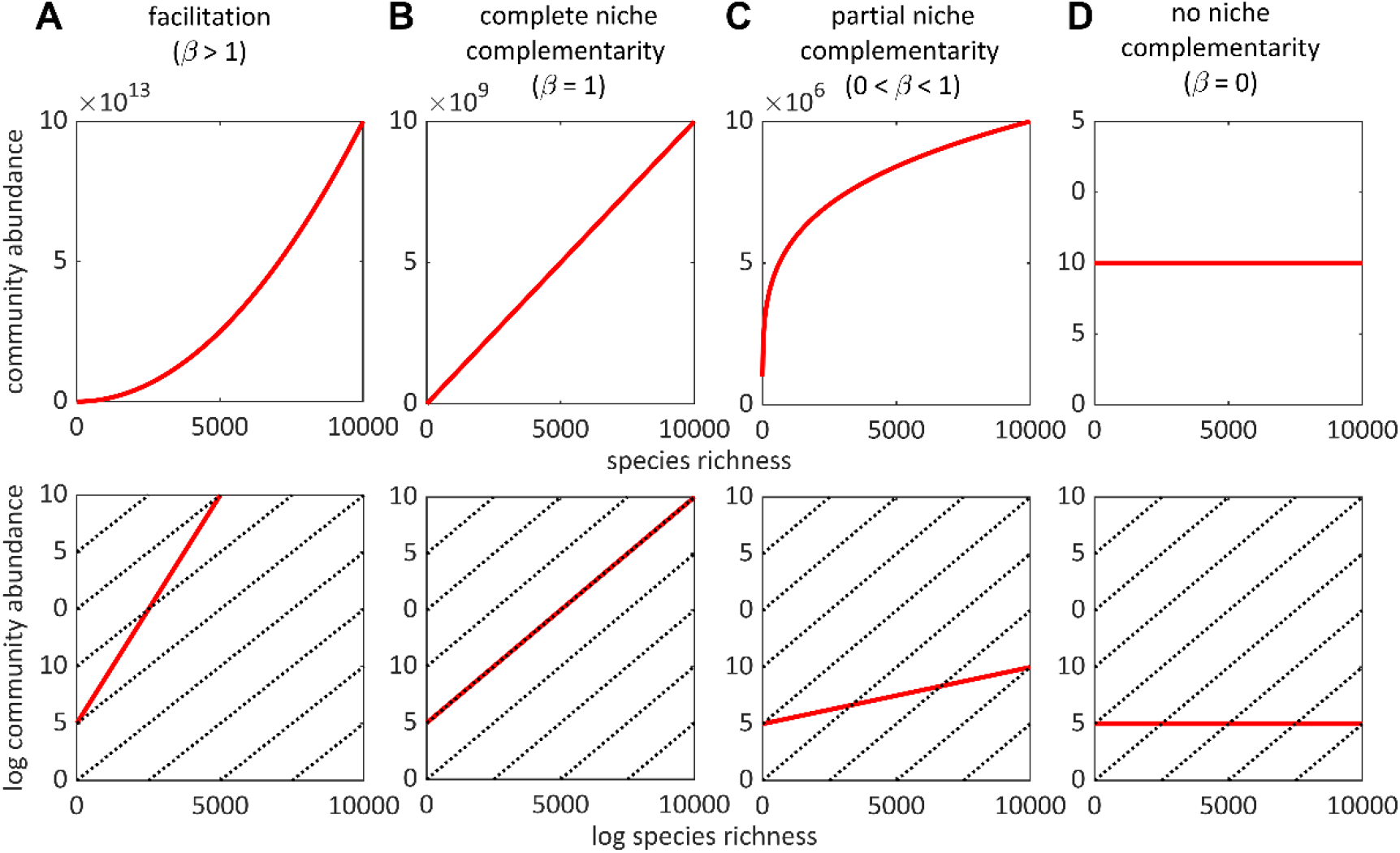
Interspecific facilitation, niche complementarity, and competition shape the J-nullcline, which reflects the biodiversity effect on equilibrium community abundance (BECA). The scaling exponent *β* is the slope of the J-nullcline in log-log space (bottom row of panels) and quantifies the strength of the BECA. Bold curves and lines illustrate different hypothetical J-nullclines and dashed lines with *β* = 1 are shown for comparison. (**A**) When facilitation pervades, adding species to a community leads to a disproportionate increase in community abundance. (**B**) If there is no facilitation but each species in the community has a unique niche (e.g., does not compete with other species for resources), then adding species to a community leads to a linear increase in community abundance. (**C**) If species have niches that are partially complementary and partially overlapping due to competing for shared resources, then adding species to a community leads to a less than proportional increase in community abundance. (**D**) If all species have identical niches, then species richness has no effect on community abundance. Note that in all scenarios, the J-nullcline plateaus at some maximum (not shown here), at which point *β* approaches zero.

In figure 1C (bottom), we show the potential scenario in which facilitation dominates the J-nullcline scaling behavior at low *S* and competition dominates the behavior at high *S*, leading to a sigmoidal J-nullcline. Such facilitation at low *S* may be prevalent in physically harsh environments, deserts, certain microbial communities (Morris et al. 2013), and other ecological systems reliant on syntrophy or ecosystem engineering (Jones et al. 1997), as primary succession is known to be driven by facilitation (Connell and Slatyer 1977; Cardinale et al. 2002; Walker and Del Moral 2003; Kjær et al. 2018; Castillo et al. 2021). However, for most environmental gradients of high-diversity contemporary communities, we expect that at large scales communities fall within a phase of the J-nullcline in which *β* is positive and less than one, (O’Connor et al. 2017).

### Stability and Emergence of Multiple Equilibria

Having shown how equilibria emerge from the intersection of S and J-nullclines and the eco-evolutionary underpinnings of the nullclines, it is necessary to elucidate the nature of the equilibria – whether they are stable or unstable. The stability of the equilibrium point [*Ŝ, Ĵ*] emerges from the interaction of the *S* and J-nullclines. The stability follows some basic rules due to differences in the timescales of births and deaths versus speciation and extinction, circumventing the need for stability analysis of a specific system of differential equations. When communities are perturbed from the J-nullcline, the absolute rate of change in community abundance is much higher than the absolute rate of change in species richness, due to the difference between ecological and evolutionary time comprising population change versus speciation/extinction, respectively. Consequently, these non-equilibrium communities follow near-vertical vectors (trajectories) until they converge on the J-nullcline, at which point their path traces the J-nullcline in the direction determined by their position relative to S-nullcline and the character of the S-nullclines (figure 1B).

Because the universal attractor S-nullcline exhibits attractor dynamics, if the J-nullcline is less steep than the attractor S-nullcline around their intersection point, exhibiting *β* < 1, then the equilibrium is stable (figure 1B). A perturbed community with *S* < *Ŝ* follow trajectories that converge on the J-nullcline and then, since the J-nullcline is higher than the S-nullcline at S < *Ŝ*, the community proceeds along the J-nullcline towards the equilibrium point. In contrast, if *β* > 1 in the vicinity of the equilibrium point, then the equilibrium point is unstable (figure 1C, bottom; see also Box 1). However, due to energetic limits to *J, J* cannot increase indefinitely as *β* > 1; eventually, at very high *J*, the J-nullcline necessarily must flatten out, leading to the J-nullcline exhibiting *β* < 1 at an upper equilibrium point.

In addition to this universal stable equilibrium point, additional lower stable equilibria may emerge in a variety of scenarios. For instance, the speciation and extinction curves could have a second intersection point, if speciation probability changes with population size as a hump-shaped function or extinction probability changes with population size as a U-shaped function (e.g. if large populations have high extinction probabilities due to their susceptibility to epidemics). This second intersection point can lead to a second S-nullcline that exhibits repulsive rather than attractor dynamics (see Box 1 for details). If the J-nullcline is sigmoidal, this can lead to an additional stable equilibrium at low S (where facilitation dominates) and an additional tipping point (Box 1).

### Scaling Relationships for Gradients in Energy, Speciation, and Extinction

ETBD highlights that there are three fundamental drivers of biodiversity equilibria: energy availability, the extinction-speciation balance, and the shape of the J-nullcline (figure 1C). We thus first address how equilibrium *S* and *J* should change as a consequence of each of these three fundamental drivers, and then present quantitative predictions for complex gradients in which more than one of these drivers are varying simultaneously.

The ETDB framework reveals that the scaling behavior of the J-nullcline, as quantified by *β*, fundamentally affects the scaling relationships between *Ŝ,Ĵ*, energy availability *E*, and average individual metabolic rate *B*. Over a given range of *S*, the J-nullcline can be mathematized as a power-law function of *S* (e.g., Liang et al. 2016), giving

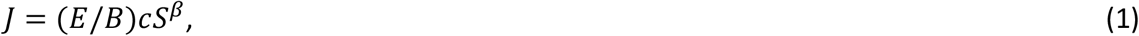

where *c* sets the fraction of the maximum available energy *E* used by a one-species community and *cS*^*β*^ ≤ 1, reflecting the physical constraint that *J* ≥ *S*. A thermodynamic constraint to Equation 1 is that *J* cannot be greater than maximum community abundance *J*_*max*_, determined by the maximum available energy *E* divided by average individual metabolic rate *B*—that is, *E*/*B* (see table 1 for a summary of ETBD’s main parameters). Equation 1 makes the baseline assumption that changes in maximum available energy *E* or *J*_*max*_ have a positive linear effect on the “height” (intercept in log-log space) of the J-nullcline (figure 1C).

**Table 1.** The main parameters of the Equilibrium Theory of Biodiversity Dynamics, its scaling theory, and model.

|  | Description | Symbol |
| --- | --- | --- |
| Core | Equilibrium number of species | $\hat{S}$ |
| | Equilibrium community abundance | $\hat{J}$ |
| | Species abundance | $N$ |
| | Equilibrium species abundance | $\hat{N}$ |
| Scaling theory | Strength of the biodiversity effect on community abundance (J-nullcline scaling exponent) | $\beta$ |
| | Maximum energy supply | $E$ |
| | Average individual metabolic rate | $B$ |
| | Environmental gradient | $G$ |
| | Exponent quantifying the effect of environmental variable $G$ on maximum community abundance ($J_{max} = E/B$) | $r_1$ |
| | Exponent quantifying the effect of environmental variable $G$ on equilibrium mean species abundance | $g_1$ |
| Model | Probability that a species of abundance $N$ goes extinct | $P_x(N)$ |
| | Probability that a species of abundance $N$ speciates | $P_v(N)$ |
| | Exponent quantifying dependence of $P_x(N)$ on $N$ | $x_1$ |
| | Exponent quantifying dependence of $P_v(N)$ on $N$ | $v_1$ |
| | Coefficient setting the extinction level at a given $N$ | $x_0$ |
| | Coefficient setting the speciation level at a given $N$ | $v_0$ |
| | Density function of the species abundance distribution | $f_N(N)$ |

Combining equation 1 with the equation for the linear S-nullcline, we obtain predictions for how *Ŝ* and *Ĵ* should change along a gradient in *J*_*max*_, *E, or B*, which is a gradient in which only the position of the J-nullcline is changing:

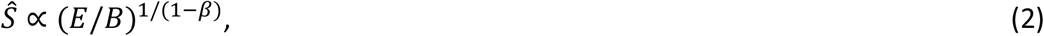

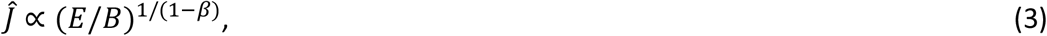

and

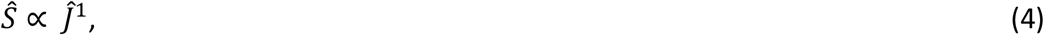

Thus, along gradients in resource availability, *Ŝ* is expected to scale linearly with *Ĵ*. In contrast, along a speciation/extinction gradient, which is a gradient in which the slope of the S-nullcline is changing, we obtain:

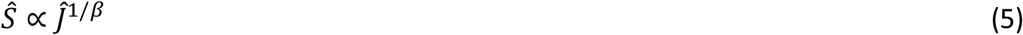

Equations 2, 3, and 5 show that the biodiversity effect on community abundance (BECA) shapes the scaling of *Ŝ, Ĵ*, and *E*/*B*. For example, when competition with niche complementarity govern the BECA (i.e., 0 < *β* <1), *Ŝ* and *Ĵ* increase superlinearly (disproportionately) with *E*/*B* and *Ŝ* scales superlinearly with *Ĵ* along speciation/extinction gradients. In contrast, when facilitation dominates (*β* > 1), *Ŝ* and *Ĵ* decrease with *E*/*B*, because with facilitation, even fewer species are required to obtain a given community abundance and equilibrium mean abundance when E/B is higher. This finding can be graphically ascertained by observing that intersection of the *S* and *J*-nullclines shifts downward and leftward when the height of the J-nullcline increases due to increasing *E*/*B (*fig. S1). Note, however, that this equilibrium point is only stable if it reflects the intersection of the J-nullcline with a second S-nullcline, which occurs when the curves quantifying the population size-dependence of extinction and speciation intersect twice (Box 1).

Equations 2-5 predict that in log-log plots of *S, J*, and *E*, straight lines are expected with slopes equal to the exponents in the equations. These equations can be taken as the most general, baseline predictions available in the absence of knowledge indicating changes in *β*. Indeed, a power law model for the BECA relationship has empirical support from studies of the effect of diversity on community biomass production, standing biomass, and resource consumption (Reich et al. 2012; Liang et al. 2016; O’Connor et al. 2017) and some theoretical basis (Mora et al. 2014; Harte et al. 2022), with *β* averaging around 0.26 across a variety of taxa and ecosystems (O’Connor et al. 2017).

We further develop the framework to quantify complex gradients along which both speciation/extinction and *E*/*B* vary, which can be used to disentangle the relative contributions of variation in speciation/extinction versus *E*/*B*. Consider the most general, coarse-grained approach: both *E*/*B* and the speciation/extinction balance (i.e., equilibrium mean abundance 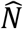) are associated with some environmental variable *G* as either power functions or exponential functions, such that

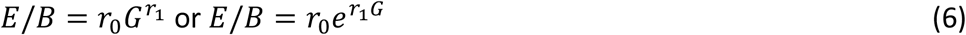

and

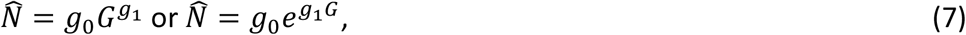

where *r*_0_ and *g*_0_ are normalization coefficients and *r*_1_ and *g*_1_ are scaling exponents quantifying the effects of *G*. The choice of whether to consider the power or exponential functions to quantify a particular gradient should be made based on theoretical or practical reasons. For instance, typically, researchers consider species richness to vary exponentially with temperature and latitude (Currie et al. 2004; Storch 2012), both because temperature has an exponential effect on biological rates (Brown et al. 2004) and because scatterplots of log richness versus temperature and latitude tend to minimize statistical heteroscedasticity compared to log-log or untransformed plots of these variables.

Along a purely resource-driven gradient, equilibrium mean abundance doesn’t vary and so *g*_1_ = 0. Along a purely speciation/extinction-driven gradient, resource availability doesn’t vary and so *r*_1_ = 0. When *g*_1_/*r*_1_ < 0, the two drivers (the positive effects of the speciation/extinction balance and resource availability on diversity) are concordant along the gradient, leading to what we call a *concordance gradient*. When *g*_1_/*r*_1_ > 0, the effects are discordant such that at higher resource availability, speciation is lower or extinction is higher, leading to a *discordance gradient*. In a perfectly discordant gradient, the drivers of diversity vary completely opposingly, leading to *g*_1_/*r*_1_ = 1 resulting in no change in diversity along the gradient.

The resulting scaling predictions for these gradients are:

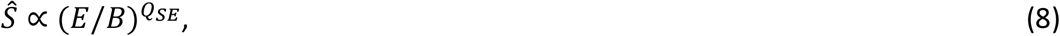

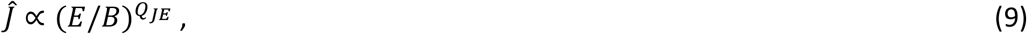

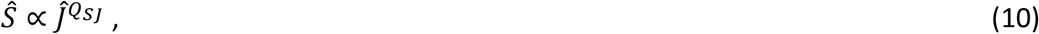

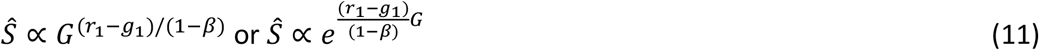

and

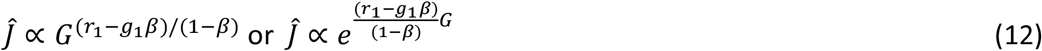

Where *Q*_*SE*_ = (1 − *g*_1_/*r*_1_)/(1 − *β*), *Q*_*JE*_ = (1 − *g*_1_*β*/*r*_1_)/(1 − *β*), and *Q*_*SJ*_ = (1 − *g*_1_/*r*_1_)/(1 − *g*_1_*β*/*r*_1_) and the formulas for Equations 11 and 12 depend on whether the gradient variable is assumed to have a power law effect (first formula) or exponential effect (second formula) on 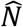 and *E*/*B*. Figure 3 and Equations 8-12 show that the scaling of *Ŝ, Ĵ*, and *E* depends only on *β* and the ratio *g*_1_/*r*_1_. For example, when *β* < 1, *Ŝ* only increases with *E*/*B* if *g*_1_/*r*_1_ < 1; that is, as long as the gradient isn’t strongly discordant. Along a discordance gradient, a great variety of *S*-*J* relationships can occur, depending on *β* and whether or not the speciation/extinction balance versus *E*/*B* is varying most along the gradient (figure 3). In a perfectly discordant gradient, although no change in biodiversity along the gradient occurs, there is a change in *J* along the gradient that is determined by *β*. Similarly to a resource gradient, along concordance gradients, *β* values higher than 1 can lead to negative S-*E* and *J*-*E* relationships, but only in the presence of a second S-nullcline. Species richness and abundance patterns along environmental/geographic gradients thus depend on (1) the degree of concordance or discordance between two major drivers of diversity variation, namely speciation/extinction rates and resource availability, and (2) the exact form of BECA.

**Figure 3.**
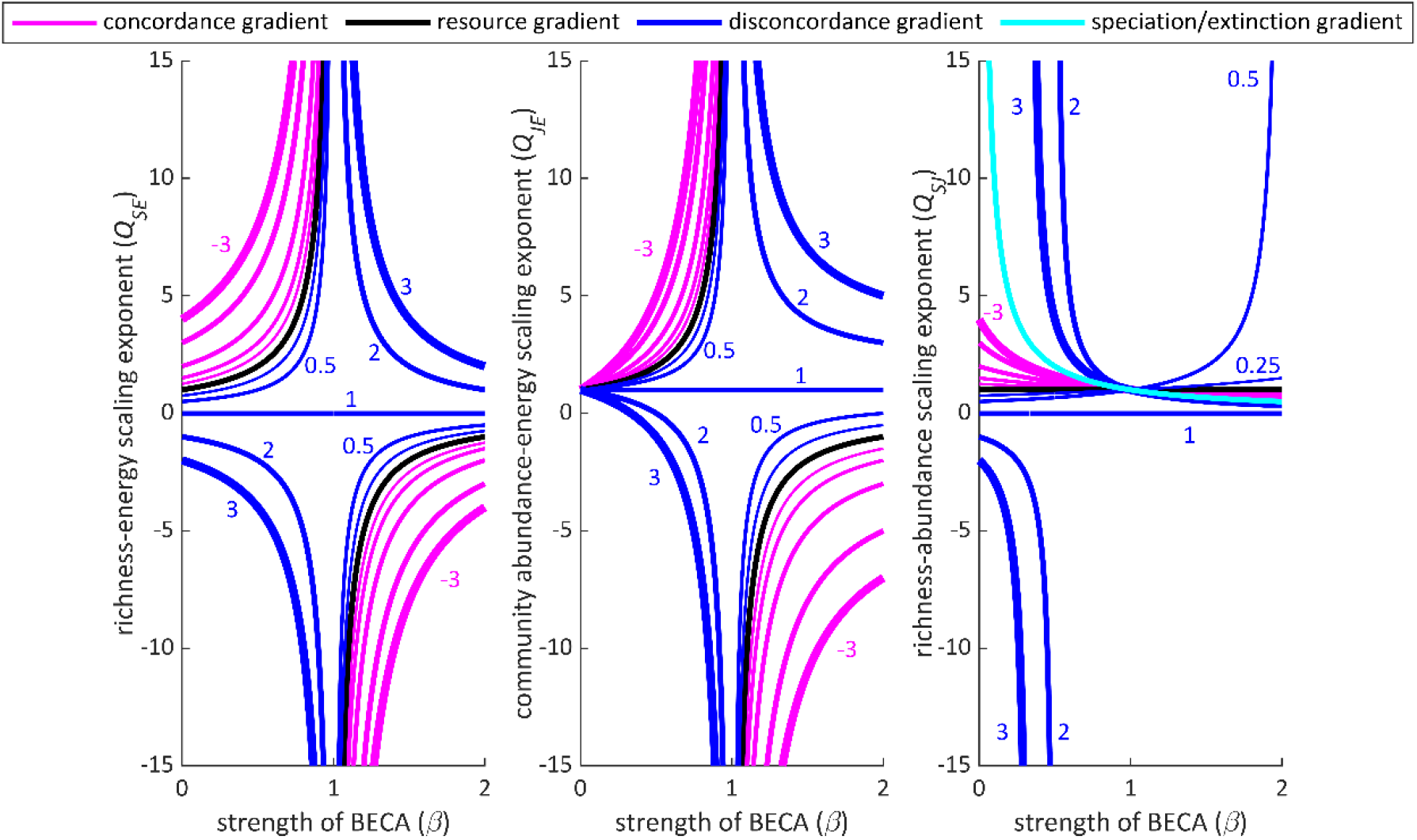
Effects of the J-nullcline on the scaling of species richness and community abundance equilibria along simple and complex environmental gradients. *β* quantifies the strength of the biodiversity effect on community abundance (BECA). The curves indicate the *S*-*J*-*E* scaling exponents for pure speciation/extinction gradients (light blue, *r*_*1*_ = 0), pure resource availability gradients (black, *g*_*1*_ = 0), and complex gradients in which the speciation/extinction balance and resource availability covary. In a complex gradient, equilibrium mean population size 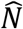 may decrease with resource availability due to higher speciation or lower extinction in high resource environments (a *concordance gradient*, pink curves, *g*_*1*_/*r*_*1*_ < 0) or increase with resources (*a discordance gradient*; dark blues curves, *g*_*1*_/*r*_*1*_ > 0). Thicker curves indicate gradients having larger absolute values of *g*_*1*_/*r*_*1*_. The value of *g*_*1*_/*r*_*1*_ is shown next to each curve. All communities have an upper equilibrium point resulting from competition for limited resources (*β* < 1) that is stable. If the J-nullcline exhibits facilitation (*β* > 1) below this equilibrium point, then communities can have additional equilibria (Box 1).

Although it may seem unintuitive, discordant gradients may occur in a variety of situations. For example, communities of microbes and ectothermic animals in some high latitude marine ecosystems may frequently have higher *E*/*B* compared to their tropical counterparts due to nutrient enrichment from upwelling zones elevating *E*, while colder temperature simultaneously leads to lower metabolic rates. In contrast, the tropical ecosystems may have lower 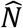, because (1) of possible positive effects of temperature on rates of genetic divergence and speciation (Allen and Gillooly 2006; Allen et al. 2006) and (2) more stable environmental conditions due to reduced seasonality and increased climatic stability that may lower rates of extinction. Therefore, in this case, the latitudinal gradient represents a complex gradient of increasing *E*/*B*, decreasing speciation, and increasing extinction with increasing latitude.

### Effects of the Species Abundance Distribution

Our framework can be further developed in order to predict how the species abundance distribution and population size dependences of extinction and speciation affect equilibrium mean abundance 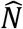 and consequently *S* and *J* equilibria. Consider communities whose species abundance distribution at equilibrium are characterized by a probability density function 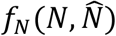 with equilibrium mean abundance 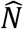, which follows from the speciation-extinction dynamics as explained above. We make the baseline assumption that all the other parameters of the probability density function, should they exist, are invariant across communities, as there is no a priori reason to assume that they change with *S* or *J*. At the beginning of a given time interval Δ*t*, each species in a community has some abundance *N* that may vary among species in the community. By the end of the interval, the proportion *P*_*x*_(*N*) of species of abundance *N* that go extinct is a function of *N*.

Likewise, some proportion *P*_*v*_(*N*) of the species of abundance *N* may speciate, with each speciation adding a new species to the community. *P*_*x*_(*N*) and *P*_*v*_(*N*) are equivalent to the probabilities that a species of abundance *N* goes extinct or speciates, respectively, within the time interval.

Integrating over the species abundance distribution, the community’s mean total number of extinctions *X*_TOT_ that occur within the time interval is 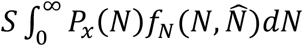 and the mean total number of speciations *V*_*TOT*_ is 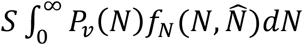. At equilibrium, the mean total speciation rate (*V*_*TOT*_/ Δ*t*) must be equal to the mean total extinction rate (*X*_*TOT*_/ Δ*t*), giving the equilibrium constraint formula

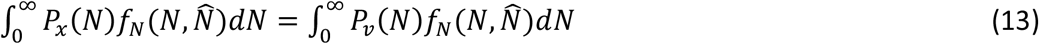

By assuming functions for *P*_*v*_(*N*), *P*_*x*_(*N*), and *f*_*N*_(*N*), equation 13 can be used to solve for equilibrium mean abundance (the slope of the S-nullcline), which in conjunction with an assumed function for the J-nullcline (characterizing the BECA), can be used to predict equilibrium *S* and *J* and their dependences on the parameters of extinction and speciation (*Materials and Methods*).

In the unrealistic, idealized scenario in which all species in the community have the same abundance (“the same-abundance distribution”), it follows from integrating Equation 13 that 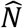 is equal to the intersection point of *P*_*x*_(*N*) and *P*_*v*_(*N*). If instead *N* varies among species (i.e., the distribution has a positive variance), then 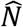 can deviate from the intersection point of *P*_*x*_(*N*) and *P*_*v*_(*N*), since 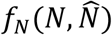 acts to weigh the degree to which species of different *N* contribute to the total rate of speciation and extinction. For example, even if species with low *N* have low probabilities of speciation, they could be important contributors to the total rate of speciation if there are many species with low *N*. Equation 13 demonstrates that the effects of the shape of the species abundance distribution on 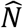 are complex, depending on its interaction with *P*_*x*_(*N*) and *P*_*v*_(*N*).

Generally, with typical species abundance distributions, such as the logseries and lognormal distribution, and ecologically-relevant functions for *P*_*x*_(*N*) and *P*_*v*_(*N*), we’ve found that increases in the distribution’s variance lead to larger values of 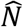 (figure 3). This is because these distributions exhibit asymmetry with many low-abundance species and few high-abundance species.

Consequently, there are many species that have higher probabilities of extinction and lower probabilities of speciation than the species with the abundance found at the intersection of *P*_*v*_(*N*) and *P*_*x*_(*N*), leading to the community as a whole needing to have a higher mean abundance in order for speciation rates to balance extinction rates. Regardless, the intersection point of *P*_*x*_(*N*) and *P*_*v*_(*N*) is the main determinant of 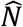 and shifts in *P*_*v*_(*N*) and *P*_*x*_(*N*) predictably move the S-nullcline and equilibrium *S* and *J* (figure 3).

### Effects of speciation and extinction and their population size-dependences

In order to provide quantitative predictions of the effects of the N-dependence of speciation and extinction rates and different species-abundance distributions, we suggest the use of *J* = (*E*/*B*)*cS*^*β*^ for the J-nullcline and derived the following baseline functions for *P*_*x*_(*N*) and *P*_*v*_(*N*) to deploy as strong starting points:

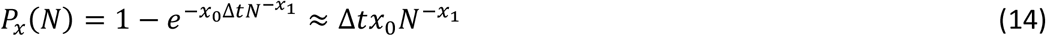

and

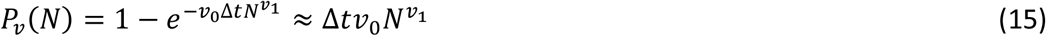

Where *x*_0_, *x*_1_, *v*_0_,and *v*_1_ are the parameters determining the rates and N-dependence of extinction and speciation, respectively, and Δ*t* is the time interval. The analytical approximations are accurate for *N* ≫ 1 (Supplementary Materials). Equation 14 is derived by building on theory for extinction driven by environmental stochasticity (Ovaskainen and Meerson 2010). The extinction scaling exponent *x*_1_ is mechanistically linked to population growth rate and its variance (Ovaskainen and Meerson 2010), and *x*_1_ ≥ 0 due to the overall negative N-dependence of extinction rate. Equations 1, 14, and 15 may be taken as operational tools to make coarse-grained predictions by quantifying the log-log slopes of the J-nullcline around equilibrium points and the slopes of *P*_*x*_(*N*) and *P*_*v*_(*N*) near their intersection points. Equation 15 thus does not discount the possibility that *P*_*v*_(*N*) is peaked nor that the J-nullcline exhibits more complex behavior (e.g., sigmoidal), as long as *β* around the equilibrium point varies negligibly along the gradient of interest. Alternatively, the J-nullcline and extinction functions may be taken as baseline models in their own right, as they have some theoretical and empirical support (see Supplementary Materials for derivations).

Assuming the same-abundance distribution and equations 14 and 15 approximations, the theory predicts how equilibrium diversity, community abundance, and mean abundance should vary along gradients in the parameters governing speciation and extinction rates:

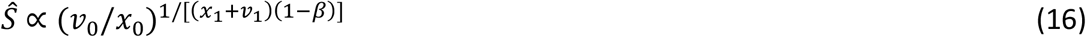

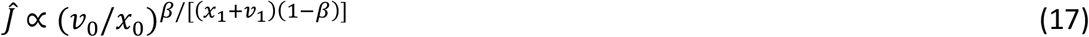

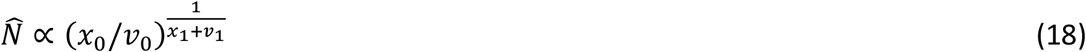

Importantly, equation 16 and 17 highlight that the effect of the levels of extinction (*x*_0_) and speciation (*v*_0_) on *Ŝ* and *Ĵ* is modulated by the biodiversity effect on community abundance (BECA) and the degree of *N*-dependence of extinction and speciation. Additionally, at equilibrium, the extinction and speciation parameters (*x*_1_, *v*_1_, *x*_0_ and *v*_0_), together with BECA and *E*/*B*, determine the total rates of extinction and speciation.

In order to evaluate the role of the species abundance distribution in modifying the diversity gradients, we determined how equilibria change along gradients in resource availability (*E*/*B*) and speciation (*v*_0_) assuming lognormal species abundance distributions with different parameters. We solved equilibria numerically, as we were unable to obtain analytical solutions. The calculations require solving for the value of 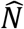 that balances the total rate of extinction with the total rate of speciation (eq. 13) and then solving for the intersection of the *S* and J-nullclines (*Materials and Methods*). Figure 4 shows that the shape of the species abundance distribution, whether the same-abundance distribution or lognormal distribution with low or high variance, has minimal impact on overall changes in biodiversity along gradients in speciation/extinction or resource availability, although it certainly modulates equilibrium mean abundance and so it could be important to consider if the shape of the species abundance distribution is suspected to change drastically over a gradient.

**Figure 4.**
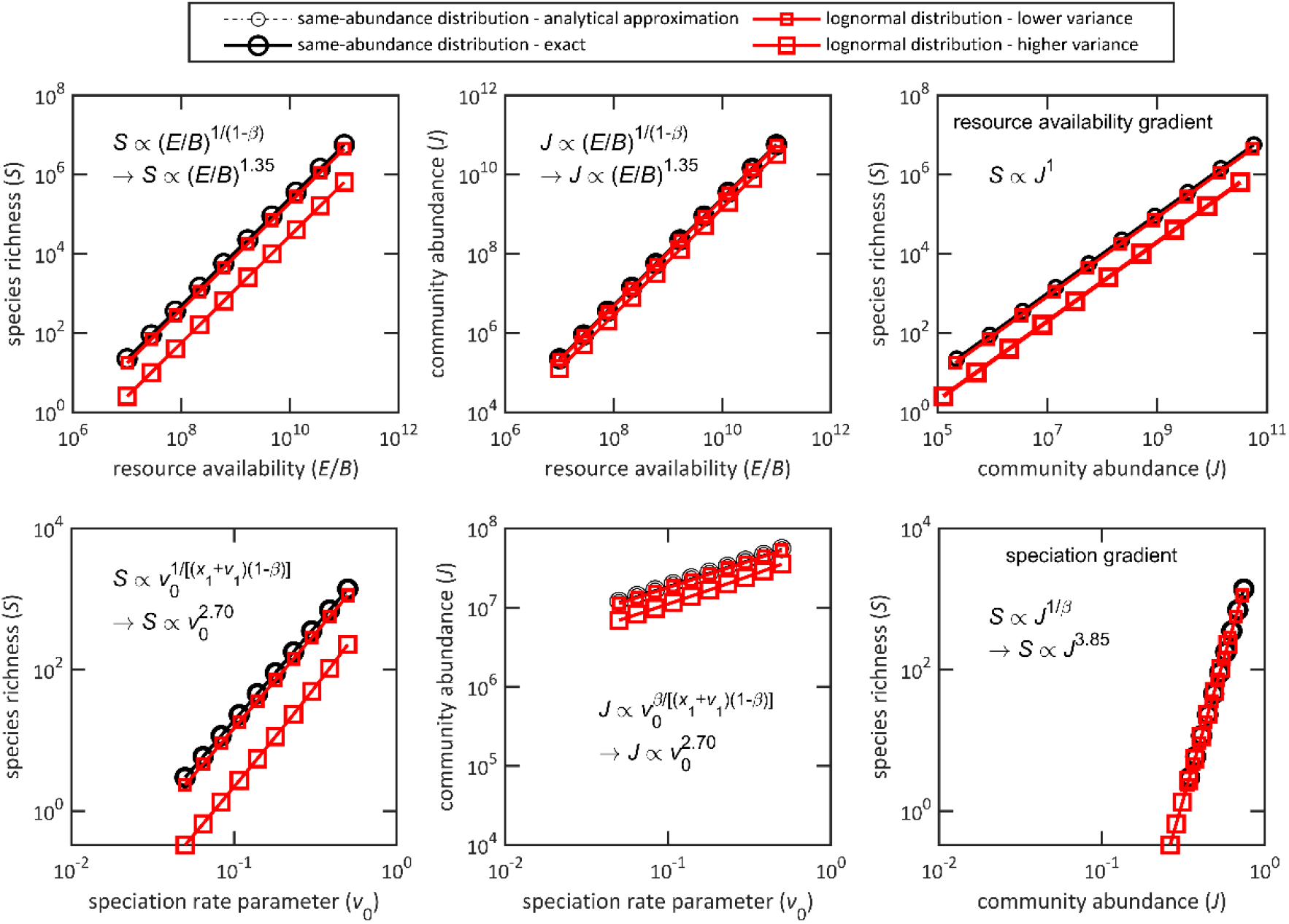
Predicted changes in biodiversity and community abundance along gradients in resource availability (top row) and speciation (bottom row) for different species abundance distributions. Symbols indicate arbitrary locations along the gradient to facilitate comparison between models. The species abundance distribution has limited effects on the scaling patterns, although it does influence the species richness and community abundance expected for a given set of environmental conditions. For communities with lognormal species abundance distributions, the distributions’ *σ* parameter (standard deviation of ln *N*) was assumed to be either 0.5 or 1.5 for low and high variance species abundance distributions, respectively. The speciation rate parameter (*v*_0_) determines the proportion of species of a given abundance that speciate within a given time interval. In all panels, *x*_1_ = 0.5, *c* = 0.01, *β* = 0.26, *v*_1_ = 0. In **A, B**, and **C**, *x*_0_ = 10, *v*_0_ = 0.1. In **D, E**, and **F**, *E*/*B* = 10^9^ individuals and *x*_0_ = 100.

### Empirical Application of the Framework

We applied the scaling framework to variation in tree species richness across the globe in order to clarify drivers of the latitudinal diversity gradient, using a multi-scale tree biodiversity database (Keil and Chase 2019)(*Materials and Methods*). Some authors argue that energy/resource availability is a major driver of latitudinal diversity gradients (e.g., Wright 1983; Gaston 2000; Storch et al. 2006; Hurlbert and Stegen 2014), whereas others have argued that variation in speciation or extinction rate is a major driver (e.g., Fuhrman et al. 2008; Brown 2014; Rolland et al. 2014; Wu et al. 2018).

Since ETBD makes alternative predictions for the *S*-*J*-*E* scaling relationships depending on the role of these factors, it can be used to shed light on the underpinnings of the latitudinal diversity gradient.

While ETBD deals with the scaling of diversity and abundance of communities representing large scales, high-quality abundance data for such large scales is rarely available. We therefore first checked whether a biome’s average local plot-level species richness scales predictably with its regional-scale (100,000 km^2^) richness, in order to determine whether the theory’s predictions have implications for across-biome variation in local richness. We found that local richness in small (0.01-0.1 km^2^) and medium-sized plots (0.1-1 km^2^) scales approximately linearly with regional richness (log-log slopes = 1.14 and 0.94, R^2^ = 0.77 and 0.75, respectively; figure 5A). In trees, it’s thus reasonable to use ETDB to elucidate the drivers of across-biome variation in plot-level richness at plot sizes down to 0.01 km^2^. This premise accords with arguments by ecologists and biogeographers emphasizing regional and biome-scale source pools as shapers of global-scale patterns of local biodiversity (Ricklefs 1987, 2004; Cornell and Harrison 2014). Note, however, that a linear relationship between local richness and regional richness is not expected to hold down to very small scales, due to the limiting effect of the low number of individuals at these scales (Storch 2016). Thus, in order for the theory to predict across-biome variation in local tree richness of plots smaller than 0.01 km^2^, the nonlinear scaling of this local richness with regional richness would have to be incorporated into the quantitative predictions.

**Figure 5.**
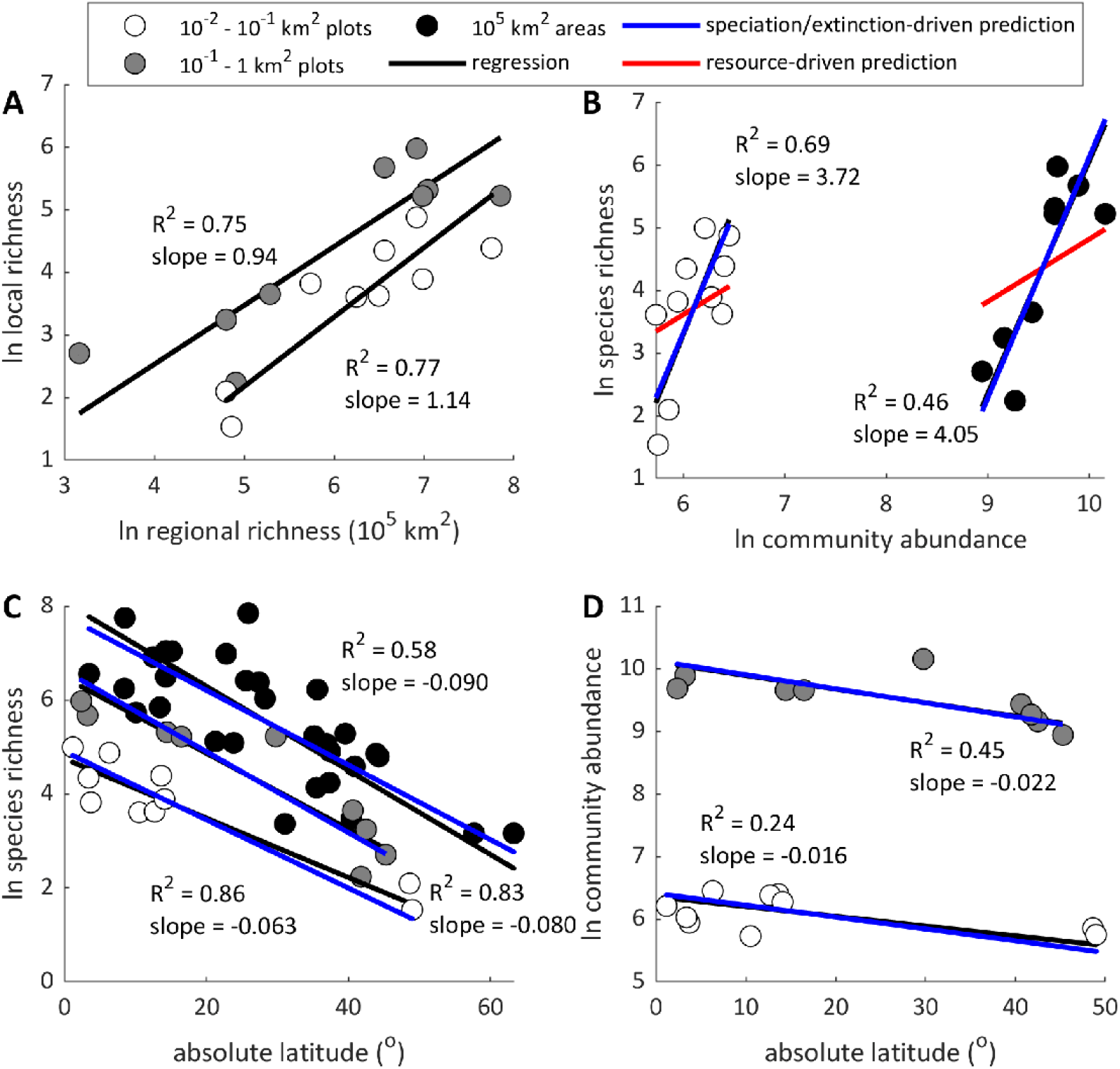
Application of the theory to among-biome patterns of average tree richness suggests the balance of speciation and extinction, rather than variation in energy availability, is the major driver of their latitudinal diversity gradient. (**A**) Local richness down to 0.01 km^2^ grain sizes is proportional to 100,000 km^2^ regional richness, indicating that in trees ETBD predictions downscale to these lower grain-sizes without need for amendment. In (**B**), (**C**), and (**D**), the blue lines show the predicted slopes for the relationships between log species richness and log community abundance, log species richness and latitude, and log community abundance and latitude assuming a purely speciation/extinction driven gradient and *β* = 0.26 for the J-nullcline. Predictions are indistinguishable from the regressions (black lines). Red lines in (**B)** represent the contrasting scenario of a purely resource-driven gradient (i.e., slope of 1).

Next, we used the theory to predict the across-biome *S*-*J* scaling relationships (*Q*_*SJ*_) for different scenarios of diversity variation and compared predictions to the observed *S*-*J* relationships to determine which scenario manifests in trees. If across-biome diversity variation is driven entirely by resource availability (*E*/*B*), then equations 4 and 9 predict *S* ∝ *J*^1^ (i.e., *Q*_*SJ*_ = 1). In contrast, if diversity variation is driven entirely by a speciation/extinction gradient, then equations 5 and 9 predict *S* ∝ *J*^1/*β*^, where *β* is the scaling exponent quantifying the biodiversity effect on community abundance (BECA). Assuming *β* = 0.26, which is suggested by an analysis of tree diversity in forests (Liang et al. 2016) and a meta-analysis of studies of BECA in primary producers (O’Connor et al. 2017), our theory predicts *S* ∝ *J*^3.85^ for this scenario. If instead diversity variation reflects a complex gradient involving a concordance of both drivers (i.e., *g*_1_/*r*_1_ < 0), then equation 9 predicts the scaling exponent to be between 1 and 3.85, whereas if there is a discordance of the drivers (i.e., *g*_1_/*r*_1_ > 0), then it predicts a scaling exponent less than 1.

For small and medium plot sizes, we found, respectively, *S* ∝ *J*^3.72^ and *S* ∝ *J*^4.06^ (R^2^ = 0.69 and 0.46), where *S* and *J* are the biome-averaged species richness and total number of individuals per plot for major terrestrial biomes in the different biogeographic realms (figure 5B). The observed average *S*-*J* scaling is thus *S* ∝ *J*^3.89^, which is remarkably close to the predicted 3.85 for a purely speciation/extinction driven gradient (i.e., where *r*_1_ = 0) whereas the 95% confidence intervals of the observed scaling relationships do not overlap with one as expected for a purely resource driven gradient (i.e., where *g*_1_ = 0). In other words, the patterns found are in accord with the idea that tree diversity variation across biomes (and thus the latitudinal diversity gradient) is not driven by variation in resource availability but rather by variation in speciation and extinction rates (possibly due to a temperature-dependence of speciation probability, see Box 2).

Assuming this scenario, we can predict the rate at which species richness and community abundance should change with latitude by using the theory’s equations. First, we estimate the value of the rate (*g*_1_) at which the speciation/extinction balance changes with latitude from the slope of log mean abundance versus absolute latitude (eq. 6), giving *g*_1_ =0.023 and *g*_1_ = 0.028 for small and medium-sized plots, respectively, where *G* is absolute latitude (table 2). Using these values of *g*_1_ we thus predict from equations 11 and 12 that *S* ∝ 10^−0.031*G*^ and *J* ∝ 10^−0.0082*G*^ in small plots and *S* ∝ 10^−0.037*G*^ and *J* ∝ 10^−0.0097*G*^ in medium-sized plots. Using the mean value *g*_1_, we also predict that at the regional-scale *S* ∝ 10^−0.035*G*^. These predictions are all remarkably close to and within the 95% Confidence Intervals of the observed (figure 5C and D, table 2).

**Table 2.**
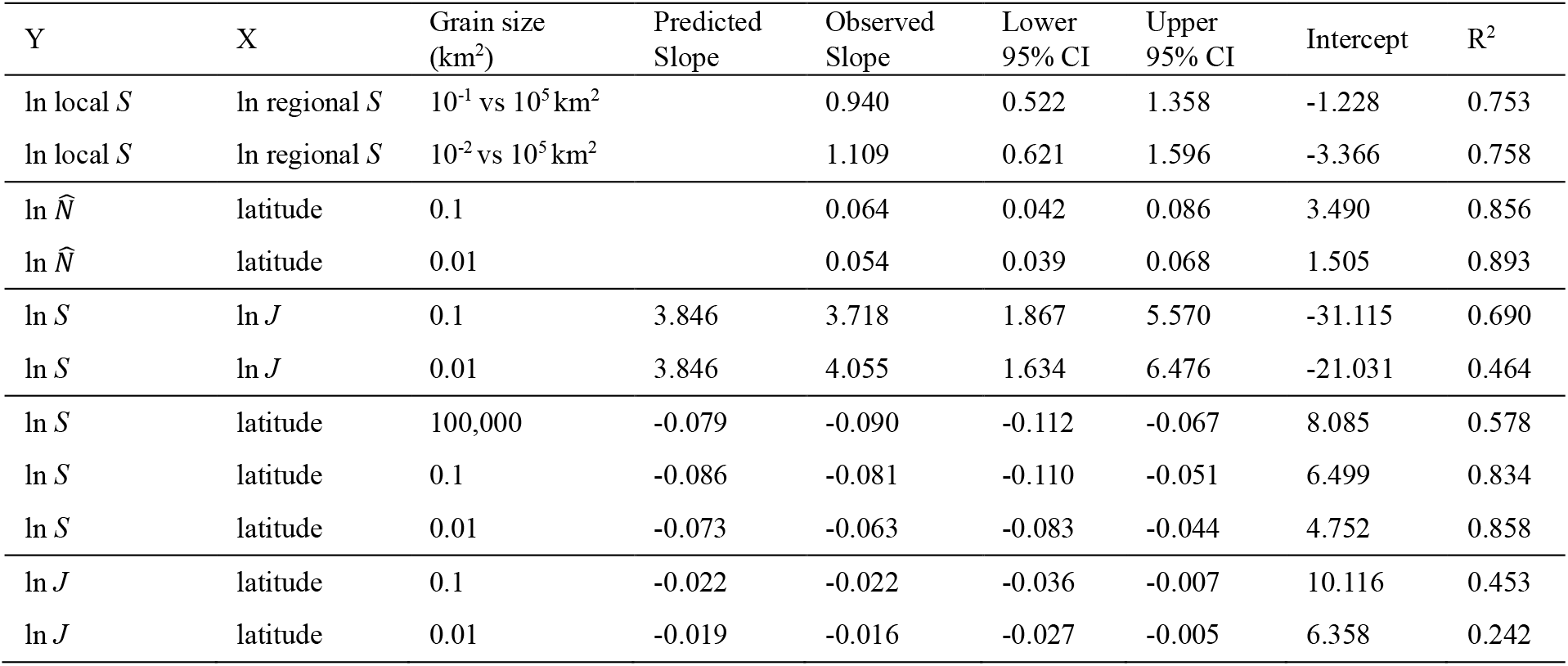
Results of the Reduced Major Axis regression analysis of the across-biome relationships between latitude, tree species richness (*S*), total number of individuals (*J*), and mean population abundance 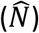, and comparisons to the ETDB-predicted slopes for a speciation/extinction-driven diversity gradient.

## Discussion

We presented general quantitative theory that predicts the scaling relationships between equilibrium species richness, community abundance, and energy availability along complex, large-scale environmental gradients and sheds light on the emergence of stable and unstable equilibria in biodiversity and community abundance. ETBD shows how nullclines for richness and community abundance emerge from a few basic assumptions and are influenced by the strength of biodiversity effect on community abundance (BECA) and the multitude of geographic, genetic and ecological factors influencing extinction, speciation, and the species abundance distribution. In doing so, the theory provides a useful unifying framework for elucidating the drivers of biodiversity patterns, such as the latitudinal diversity gradient. While we develop general scaling theory and a baseline model involving a minimal number of assumptions, more complex scenarios of eco-evolutionary dynamics and complex gradients can be incorporated into the framework in order to address particular interactions between ecological and evolutionary processes.

### Theoretical Overview

ETBD reveals shortcomings of existing biodiversity theories for explaining general patterns of diversity, including the original metabolic theory of ecology (Box 2), the MacArthur-Wilson theory of insular biogeography (MacArthur and Wilson 1967), the Neutral Theory of Biodiversity (Hubbell 2001), and their descendants (Worm & Tittensor 2018). No previous general theory of biodiversity—one that predicts biodiversity as a function of abiotic and biotic variables—has addressed the complex interactions between species richness and community abundance, namely the positive effect of species richness on community abundance.

Our framework unifies theories on the role of community abundance in diversity regulation due to mechanisms of “the More Individuals Hypothesis”, which predicts a linear effect of community abundance on species richness (Storch et al. 2018), with work on the biodiversity-ecosystem functioning relationship (Loreau 1998; Cardinale et al. 2006; Mora et al. 2014; Liang et al. 2015; O’Connor et al. 2017; Xu et al. 2020). We found that the resulting feedbacks between biodiversity and community abundance have fundamental repercussions for the stability, scaling, and gradients of equilibria, with the strength of the biodiversity effect on community abundance (BECA) mediating the effects of energy availability on species richness (*S*) and community abundance (*J*), as well as the relationship between species richness and community abundance along gradients in per species extinction or speciation rates. Thus, for the purposes of macroecological predictions, community abundance cannot be taken as an independent static variable, as in prevailing biodiversity theory and models (MacArthur and Wilson 1967; e.g., Hubbell 2001; Brown et al. 2004; Harte 2011). Indeed, statistical analyses of geographic variation in biodiversity and community biomass/productivity, employing approaches such as structural equation models, suggest bi-causal relationships between species richness and community biomass/productivity, corroborating this point (e.g., Grace et al. 2016; Craven et al. 2020). An important finding resulting from our integration of these phenomena is that a positive biodiversity effect on community abundance should lead to superlinear scaling between *S* and *J*, as well as between *S* and energy availability, in a variety of commonplace scenarios in communities experiencing competition combined with niche complementarity. This is in accord with previously enigmatic observations of superlinear scaling (Brown 2014; Storch et al. 2018; Hamilton et al. 2020).

Importantly, ETBD shows that all persistent communities (communities that aren’t on a path to extinction) have at least one stable equilibrium in biodiversity and community abundance. This stable equilibrium emerges from two universal constraints to biodiversity dynamics—that (1) the curves quantifying the population size–dependence of extinction and speciation must necessarily intersect and (2) the biodiversity effect on community abundance is ultimately constrained by thermodynamic limits to community abundance. Even when communities aren’t at equilibrium, these attractors can shape biodiversity patterns in two ways. First, many communities appear to hover around equilibrium points, as suggested by accumulating evidence from macroecological and macroevolutionary analyses (Rabosky 2009, 2022; Rabosky and Hurlbert 2015; Storch and Okie 2019; Rineau et al. 2022; Šímová et al. 2023), which would lead to their average species richness being close to their equilibrium points. Secondly, following a major global-scale perturbation that pushes communities out of equilibrium, all-else-being-equal, the non-equilibrium communities with higher stable equilibria are more likely to have higher species richness than the non-equilibrium communities with lower stable equilibria, both because these communities have a greater “distance” to fall from and because the further a community is from a stable equilibrium point, the greater its rate of increase in diversity, leading to the high-equilibria community bouncing back to a higher richness more quickly. While these points may seem trivial, they can help resolve longstanding debates regarding the role of “cradles”, “museums”, and “graves” in shaping global-scale biodiversity patterns (Vasconcelos et al. 2022).

### The Latitudinal Diversity Gradient in Trees

Application of our framework to global-scale variation in tree biodiversity suggests that tree diversity variation is driven by variation in the speciation and extinction probabilities whose effects are modulated by the biodiversity-ecosystem functioning relationship, whereas energy availability (*E*/*B*, i.e., maximum attainable community abundance) plays a negligible role. We suspect three non-mutually exclusive explanations for the surprising finding of this apparent negligible role of energy/resource availability. First, maximum attainable community abundance (per unit area) may be strongly limited by space in trees (e.g., as implied by Enquist et al. 2009; West et al. 2009), which would not vary across biomes, and the size of trees, which does not vary strongly with latitude (Stegen et al. 2011). Second, any variation in maximum energy supply (*E*) across biomes may be balanced by co-variation in individual metabolic rate (*B*), with trees having higher metabolic rates in higher energy environments, such that maximum attainable community abundance (*E*/*B*) varies little across biomes. The prediction and observation that forest stand biomass is relatively invariant of mean annual temperature and latitude corroborates this point (Enquist and Niklas 2001; Stegen et al. 2011). Third, per species speciation and/or extinction rates vary substantially across biomes due to possible temperature-driven differences in rates of genetic divergence as well as differences in seasonality, climatic fluctuations from Glacial-Interglacial Cycles, and other factors.

Our finding may seem to contrast with work emphasizing the role of ecosystem productivity (such as NPP) in generating macroscale biodiversity patterns (e.g., Bohdalková et al. 2021), but it does not exclude a role for total energy or resource supply (*E*) in biodiversity regulation. Energy availability (in terms of the resource inflow) may play a more important role in biodiversity patterns of consumers, especially endotherms whose individual metabolic rates co-vary little with ecosystem production. More importantly, energy availability, according to the theory, always plays a crucial role in stabilizing diversity dynamics even when it does not directly drive spatial diversity variation.

### The Biodiversity Effect on Community Abundance (BECA)

In our analysis of tree diversity, the scaling of *S* with *J* was remarkably well-predicted assuming the exponent *β* of 0.26 for the BECA. This support for the theory’s predictions with 0 < *β* < 1 suggests these biomes hover around their upper tree diversity equilibrium points at which partially overlapping niches and competition mediate the biodiversity effect on community abundance, as opposed to a possible lower stable equilibrium point created by *β* > 1 where the effects of interspecific facilitation would dominate. Since *β* is far from zero, many communities with relatively lower species richness are far from their absolute maximum community abundances imposed by thermodynamic limits. This corroborates analyses of a variety of manipulative biodiversity experiments underscoring the pervasiveness of a positive biodiversity effects (Tilman et al. 2001; Bell et al. 2005; e.g., Cardinale et al. 2006; O’Connor et al. 2017; Qiu and Cardinale 2020), with the majority of exponents quantifying the effects of species richness on community biomass, productivity, and resource availability being between 0 and 1, averaging 0.26, (O’Connor et al. 2017) and reported to be 0.26 for the effect of richness on tree community productivity in forests (Liang et al. 2016).

In plants, the positive biodiversity effect on community abundance can be expected, among other things, to result from the differing effects of species in modulating biomass loss due to herbivory and diseases, the role that different species play in re-shaping nutrient limitations to growth, and other factors related to niche efficiency and complementarity (e.g., Loreau 1998; Liang et al. 2015; Wright et al. 2017). In consumer communities, different species can vary substantially in their ability to acquire and metabolize different resources. Many resources go unused by animals, contributing instead to decomposer pools, and the introduction of an animal with an innovation in foraging or digestion can improve the acquisition or assimilation of a resource (e.g., DeMiguel et al. 2014; Pyenson 2017), increasing the fraction of total energy that is converted into community abundances.

Our theory and analyses underscore a need to better understand the fundamental quantitative features of the biodiversity effect on community abundance. Although the scaling exponent of the J-nullcline could depend on spatial scale (Gonzalez et al. 2020; Barry et al. 2021), the actual extent of this scale-dependence is unknown and a positive effect of diversity is expected to hold across scales (Gonzalez et al. 2020; Qiu and Cardinale 2020). Throughout Earth history, biological diversification has repeatedly expanded the total energy budget and biomass of biomes and the biosphere (Kennedy et al. 2006; David and Alm 2011; Erwin et al. 2011), indicating the effects of biodiversity on community abundance at scales of biomes and the Earth system.

Determining the strength of this biodiversity effect at these large scales is an important but challenging avenue for future research. Addressing the severe gaps in contemporary and paleontological abundance data, particularly for entire communities or assemblages (Dirzo et al. 2014), is crucial for this endeavor and for using ETBD to evaluate the role of community abundance and resource availability in shaping patterns and trajectories of biodiversity.

### Emergence of multiple biodiversity equilibria

While the role of interspecific interactions in creating multiple equilibria in community structure or ecosystem properties has been previously recognized (e.g., Holling 1973; Sutherland 1974; Scheffer et al. 2001; Konar and Estes 2003), we have outlined a general theory that quantifies the effects of niches, competition, and facilitation, as well as nonlinearities in the population size-dependence of extinction and speciation, in creating multiple equilibria in biodiversity and community abundance ’(Box 1). As such, ETBD provides a formal quantitative framework for informing qualitative arguments over the existence of planetary-scale tipping points (e.g., Brook et al. 2013; Lenton and Williams 2013) and the role of facilitation and ecosystem engineering in shaping historical patterns of diversification and biosphere modification (Jones et al. 1994). Our theory demonstrates how planetary-scale tipping points could emerge more readily than previously thought (Brook et al. 2013). In any case, just as many ecosystems appear to have multiple stable states in key ecosystem properties (Scheffer et al. 2001; Kéfi et al. 2016), ETBD suggests that many local and biome-scale ecological systems could have multiple stable biodiversity equilibria. Consideration of such multiple stable equilibria and critical thresholds that emerge from facilitation may be vital towards understanding some historical and geographic shifts in biodiversity and biomass, as well as biodiversity responses to human-induced changes in the Anthropocene (Storch et al. 2022).

### Concluding Remarks

We have presented a theory that integrates major processes underlying diversity dynamics into a single predictive framework. A strength of the framework is that it combines generality with flexibility. The theory concerning the scaling of species richness and community abundance along environmental gradients makes baseline predictions that do not require specific assumptions about ecological or evolutionary mechanisms, while specific mathematical models can also be implemented to address particular scenarios of evolution, community assembly, or global change. For example, under neutral community assembly, the species abundance distribution is a logseries distribution, whereas under purely niche-based assembly, the species abundance distribution tends towards a lognormal distribution (Pueyo et al. 2007). ETBD can accommodate either extreme of assembly, as well as the niche-neutral continuum, by assuming one of these species abundance distributions or a generalized species abundance distribution quantifying the entire assembly continuum (Pueyo et al. 2007). The theory can also be used to evaluate specific scenarios in which assembly, ecosystem functioning, speciation, or extinction are hypothesized to change along an environmental gradient or with global change (e.g., Box 2), pointing to future directions for developing a predictive ecology of biodiversity change in the Anthropocene. While ETBD is not a substitute for processed-based simulation models of biodiversity dynamics (Rangel et al. 2018; Pontarp et al. 2019; Hagen 2023), it demonstrates the continued usefulness of mathematical theory in macroecology and points to crucial eco-evolutionary phenomena that should be addressed in detailed biodiversity models.

## Materials and Methods

### Solving for Equilibrium S and J

In order to predict how *S* and *J* should change as a function of parameters of extinction, speciation, and the species abundance distribution, specific functions need to be assumed or derived for *P*_*x*_(*N*), *P*_*v*_(*N*), and the species abundance distribution. Assuming a lognormal species abundance distribution and the functions we derived for extinction and speciation (eq. 14 & 15, Supporting Online Material), the integration of the equilibrium constraint formula (eq. 13) cannot be solved analytically. We used MATLAB 2021a (Mathworks 2021) and its function *integral* to numerically solve the integrals for equilibrium mean abundance 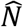–that is, for the slope of the S-nullcline. *Ŝ* and *Ĵ* were then calculated algebraically by solving for the intersection of the S and J-nullclines–that is, 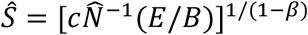 and 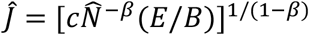. As the lognormal distribution is a two-parameter distribution (*σ* and *μ*), a value for one of its parameters must be assumed in order to solve for the other parameter and thus 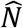 (since in a lognormal distribution 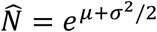). As *μ* represents the mean of log *N* whereas *σ* represents the standard deviation of log *N*, the baseline theoretical expectation is to attribute variation in 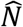 along a gradient to variation in *μ* and so assume that *σ* does not change along the gradients. Equation 13 is used to solve for *μ* and then 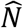 can be calculated from the relation between mean and *μ* in the lognormal distribution (i.e., from 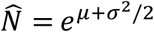.

### Data & Analysis

We used a global tree species richness database produced by Keil et al. (Keil and Chase 2019). The data include the species richness and total number of individual stems counted in plots of different sizes ranging from 0.0001 km^2^ to 0.5 km^2^, estimates of tree species richness and total number of individual trees for countries of the world and the States of the USA, and estimates of mean annual air temperature of plots and countries. Additionally, Keil and Chase (2019) categorized the biome and biogeographic realm that each plot and country is located in.

Before analyzing the data within our framework, we further curated and standardized the data in the following ways in order to increase its appropriateness for our macroecological analysis. To remove the confounding effects of isolation on biodiversity, we removed all plots and countries located on islands from our analysis. A few plots were in environments highly disturbed by humans (e.g., clear-cut forests) and so they also have been removed. The minimum diameter at base height (DBH) of trees varied some between studies, but the majority had minimum DBHs of 10 cm. To mitigate the contribution of the variation in DBH as a confounding factor, we only included plots with a minimum DBH of 10 cm.

### Country/state-level data

We used the country/state level data to estimate the average regional-scale species richness of each biome-realm combination available in the database (e.g., “Tropical and Subtropical Moist Broadleaf Forests - Indo-Malay” and “Tropical and Subtropical Moist Broadleaf Forests - Afrotropic”). Specifically, we estimated the average number of species per 100,000 km^2^ for each biome-realm combination in the following way. For each biome-realm combination, we performed an ordinary least-squares regression analysis of the log of country/state species richness versus the log of country/state area for all the countries/states found in that biome-realm combination using Minitab Statistical Software (version 2023). We then used each biome-realm combination’s parameterized regression model to estimate the number of species per 100,000 km^2^ for the biome-realm combination (i.e., regional species richness). This approach normalizes for variation in the areas of countries, thereby removing area as a confounding variable. Table S3).

### Plot-level data

We binned the plot data according to plot area, using bin widths of one order of magnitude. In practice most of the bins had much lower variation in plot sizes, as the bins were mostly made up of plots of similar standard areas (and minimum DBH). The bin for plots of 0.01 to less than 0.1 km^2^ was almost entirely made up of plots of 0.01 km^2^, so we only included plots of 0.01 km^2^ for this bin, in order to reduce confounding error. The bin for plots of 0.1 to less than 1 km^2^ had plot areas varying from 0.14 km^2^ to 0.5 km^2^ with the majority having sizes of 0.2-0.25 km^2^. For each plot area bin, we calculated the mean of the following variables for each biome-realm combination: log species richness, log number of individuals, mean annual air temperature, and mean absolute latitude (Table S3).

### Diversity gradient analysis

We used Reduced Major Axis regression to determine the relationships between variables, log-transforming variables when appropriate in order to linearize the expected relationship between variables. We used MATLAB R2021a and the formulas from Niklas (Niklas 1994) to calculate parameter values and their 95% Confidence Intervals. Reduced Major Axis regression was chosen because it is a Type II regression model (Sokal and Rohlf 1981) employed for estimating the parameter values of functional, structural, scaling, or law-like relationships between variables (McArdle 1988; Smith 2009). To evaluate the sensitivity of our results to our method for calculating regional species richness, we also repeated all regional species richness analyses (local versus regional richness and regional richness versus absolute latitude) using raw country/state richness data without implementing the normalization for variation in country/state area. In other words, for each biome-realm combination, we calculated the mean log country/state species richness and repeated the previously described local-regional scaling and gradient analyses on this measure of regional richness. The results from this analysis were similar to our other findings (Table S1).

## Acknowledgements

We thank Drew Allen, Fangliang He, Levi Gray, Robert Pacák, Honza Smyčka, Anna Toszogyova, the Storch research group, and the Network for Ecological Theory Integration (NETI) for helpful discussions and support; Dan Rabosky, Allen Hurlbert, and an anonymous reviewer for valuable feedback on the manuscript; the Czech Science Foundation (20-29554X), National Aeronautics and Space Administration (NNX16AJ61G), and an International Mobility Fellowship from Charles University for funding. We have no competing interests to declare.

## Author contributions

JO and DS conceived the theory; JO developed the mathematical framework and models; JO and DS contributed to visualization; JO analyzed the data; DS provided feedback on theory development and analysis; JO wrote the first draft; JO and DS revised the draft.

## Data and materials availability

All data are available at Keil and Chase (2019). Code and processed data supporting the results have been archived in citable format at Zenodo by Okie and Storch (2024; https://doi.org/10.5281/zenodo.13207797).

## Box 1.

Emergence of multiple diversity equilibria and tipping points.

When the speciation and extinction curves (i.e. the dependencies of the speciation and extinction probabilities on population size) have a second intersection point, due to a hump-shaped speciation curve or U-shaped extinction curve (Box figure 1), an additional S-nullcline emerges that can lead to additional stable and unstable equilibria. A hump-shape in the speciation curve may arise due to a variety of genetic, geographic, ecological, and evolutionary factors associated with species abundance that could cause a drop in speciation rate in highly abundant species. A U-shaped extinction curve may arise if large populations have high extinction probabilities due to their susceptibility to epidemics. In such scenarios, the extinction curve slope is greater than the speciation curve slope at the second intersection, leading to the second S-nullcline exhibiting *repulsive* dynamics, such that d*S*/d*t* < 0 for communities to the left of this *repulsive S-nullcline* (blue region in Box figure 1B) and d*S*/d*t* > 0 for communities to the right of the repulsive S-nullcline (pink region). Consequently, as shown in Box figure 1B, if *β* < 1 at the *J*-nullcline’s intersection with a repulsive S-nullcline, the resulting equilibrium point is unstable, whereas the equilibrium point is stable if *β* > 1. When the *J*-nullcline is sigmoidal, the presence of two S-nullclines can thus create a lower stable equilibrium in the accelerating phase of the J-nullcline (where *β* > 1) and an unstable equilibrium point (threshold) separating this lower stable equilibrium from the universal upper stable equilibrium (where *β* < 1). Tipping points leading to state shifts and hysteresis can occur due to temporary perturbations to the nullclines kicking a community temporarily out of equilibrium into another basin of attraction, the community subsequently being stuck in the alternative stable state even upon returning to initial conditions. Such tipping points could also occur under incremental changes in the height of the *S*-nullclines or *J*-nullcline. Here the system stays in equilibrium up until the bifurcation point, where a small change in conditions shifts the biome to the alternative stable state.

**Box 1 figure 1.**
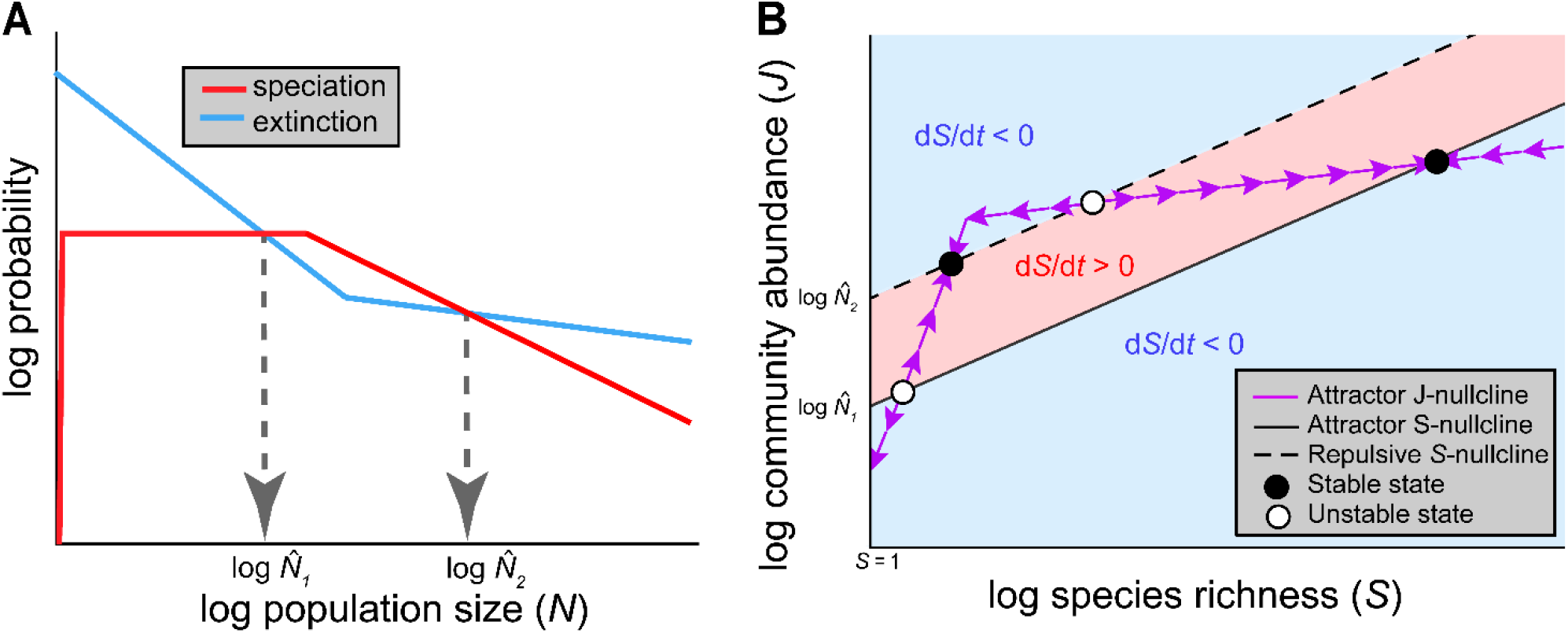
Example of how a sigmoidal biodiversity effect on community abundance (J-nullcline) in conjunction with nonlinearities in the speciation or extinction curve lead to multiple stable states in biodiversity and community abundance. (**A**) When there are two intersections of the curves, there can be two mean population sizes (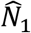 and 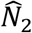) for which a community can be at equilibrium. (**B**) These equilibrium mean population sizes determine the heights (intercepts) of the resulting S-nullclines in the log-log S-J phase plane (black lines). A community recovering from a perturbation follows a relatively vertical trajectory to the J-nullcline (purple line) and then follows a trajectory along the J-nullcline (purple arrows). A sigmoidal J-nullcline is shown (like in fig 1C but in log-transformed space), leading to four equilibrium points. In **A**, the speciation curve necessarily drops to zero at non-reproductively viable population sizes, whereas the extinction curve intercepts 1.

## Box 2.

Integrating the Equilibrium Theory of Biodiversity Dynamics (ETBD) with the metabolic theory of ecology to predict diversity and abundance along temperature gradients in ectotherms, plants, and microbes.

According to metabolic theory, speciation rate should scale as 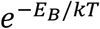 due to metabolic rate governing the temperature dependence of genetic divergence rates, giving 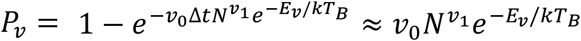, resource supply rate should scale with temperature as 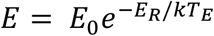, and metabolic rate should scale as 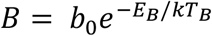, where *E*_*v*_, *E*_*R*_, *E*_*B*_ are the kinetic parameters (effective activation energies) quantifying the temperature dependence of the rates, *k* is Boltzmann’s constant in electron volts, *v*_0_, *E*_0_, and *b*_0_ are normalization coefficients, and *T*_*B*_ and *T*_*E*_ are body and ecosystem temperatures, respectively, in kelvin. Metabolic theory assumes *E*_*v*_ = *E*_*B*_ and *T*_*B*_ ≈ *T*_*E*_ in ectotherms. Both MTE and neutral biodiversity theory implicitly assume *v*_1_ ≈ 1, as they model speciation as a per capita process (Allen et al. 2006; Allen and Savage 2007). We do not make this strong assumption, as it has limited theoretical or empirical support and models of speciation suggest the population size-dependence is more complex in real populations (Gavrilets 2003). Thus, assuming the same-abundance species abundance distribution we obtain 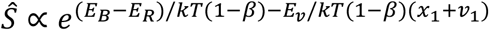 and 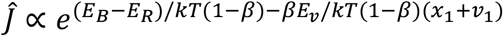. Adopting metabolic theory’s previously implicit assumption that *E*_*R*_ = *E*_*B*_, such that maximum community abundance (*E*/*B*) is independent of temperature, we obtain

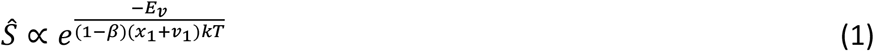

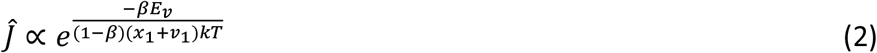

and

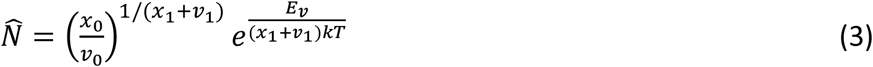

These equations show that the population size-dependence of extinction and speciation, along with the biodiversity effect on community abundance (BECA, quantified by *β*), affect the temperature dependence of diversity. This contrasts with metabolic theory’s prediction that *J* is invariant of temperature and *S* ∝ *e*^−0.65/*kT*^ due to the temperature dependence of richness in plants and ectotherms solely reflecting the temperature dependence of rates of speciation or interspecific interactions (Allen et al. 2002; Brown et al. 2004). Metabolic theory’s predictions fall well outside the 95% Confidence Intervals in trees (Table S2). Using equation 3 and assuming *E*_*v*_ = 0.65 eV (Allen et al. 2006), we calculate from our plot-level tree analysis that on average *x*_1_ + *v*_1_ = 0.70 in trees, leading to the predictions from equations 1 and 2 of S ∝ *e*^−1.26/*kT*^ and *J* ∝ *e*^−0.33/*kT*^ which are remarkably close to the observed diversity-temperature scaling relationships (Table S2). This is in agreement with the postulate of metabolic theory that the probability of speciation for populations of a given population size positively and predictably depends on temperature. However, the dependence of species richness and community abundance on temperature is only accurately predicted after accounting for the biodiversity effect on community abundance and the effects of the population size-dependence of extinction and speciation, pointing to the usefulness of ETBD in the study of diversity gradients.

