## Supplemental Online Material for "The Equilibrium Theory of Biodiversity Dynamics: a general framework for scaling species richness and community abundance along environmental gradients"

**This file includes:**

Supplementary Text
*Emergence and Properties of the S-nullcline*
*Notes on the species abundance distribution*
*On general features of* $P_v(N)$ *and* $P_x(N)$
*Derivations of* $P_v(N)$ *and* $P_x(N)$
*References*
Fig. S1
Fig. S2
Table S1
Table S2
Table S3

**Supplementary Text**

*Emergence and Properties of the S-nullcline*

Consider that at equilibrium, the total rate of extinction $X_{TOT} = S\int_0^\infty P_x(N) f_N(N,\hat{N})dN$ must be equal to the total rate of speciation $V_{TOT} = S\int_0^\infty P_v(N) f_N(N,\hat{N})dN$, giving

$$\int_0^\infty P_x(N) f_N(N,\hat{N})dN = \int_0^\infty P_v(N) f_N(N,\hat{N})dN \qquad \text{(S1)}$$

Where $P_x(N)$ and $P_v(N)$ are the functions characterizing the dependence of extinction and speciation probability, respectively, on species abundance *N* (expressed in the same units as community abundance), $f_N(N,\hat{N})$ is the probability density function of the species abundance distribution, and $\hat{N}$ is equilibrium mean species abundance.

A feature of the species abundance distribution (SAD) is that it must have a mathematically defined mean species abundance, and as such $f_N(N)$ can itself be considered a function of mean abundance. Often a probability density function has a parameter directly equivalent or related to the mean (as in the normal distribution), simplifying the mathematics, but it is not a requirement of the theory. Since $\hat{N}= J/S$, we write the SAD probability density function as $f_N(N,\hat{N})$, highlighting the fact that the equilibrium distribution is necessarily tied to the ratio of *J* to *S*. Put another way, we assume the SAD has one parameter that affects $\hat{N}$, the value of which reflects the ratio of *J* to *S* at equilibrium, which, as we shall see, is determined by the balance of extinction and speciation modulated by the SAD. The SAD may also have shape or scale parameters required to uniquely define the distribution, such as the parameter $\sigma$ in the lognormal distribution, which is typically associated with the variance. Such additional parameters, should they exist, are assumed here to be intrinsic properties of the assembly dynamics and so input parameters to ETDB. For baseline theory and modeling, we make the null assumption that all such input parameters, should they exist, are independent of *S* and *J*. Thus, we have made the assumption that the type of probability distribution characterizing the SAD (i.e., the mathematical function for the distribution) is independent of $S$, $J$, and the extinction and speciation parameters. So, for example, if the SAD is characterized by a probability distribution with two parameters, such as the lognormal distribution, then we make the baseline assumptions that: (1) all the SADs have a lognormal distribution, (2) one of the parameters is an input parameter that is invariant across communities, and (3) the remaining parameter, from which mean abundance is calculated, is determined by solving for the value that produces a diversity equilibrium (solving for $\widehat{N}$ in equation S1). As such, this parameter, along with mean abundance, may differ between communities if the parameters of extinction or speciation differ between communities. If the SAD is characterized by a one-parameter probability distribution, such as the same-abundance distribution, geometric distribution, or Fisher's log-series distribution, then there are no input parameters to assume and the properties of the SAD's probility distribution are fully determined by the extinction and speciation dynamics determining $\widehat{N}$ at equilibrium.

It is worth noting that the frequency distributions of the species abundances (such as frequency histograms produced by field biologists) are not expected to be invariant across communities differing in $S$ and $J$. The frequency distribution of the species abundances for a sample of a community characterized by a particular probability density function (e.g., as indicated by a histogram) is statistically dependent on the number of species and total abundance in that sample. For example, the number of species within an abundance class or bin increases with the number of species sampled, and the abundance of the most abundant species generally increases with the total number of individuals in the sample.

Equation S1 implies that $\widehat{N}$ is some implicit or explicit function $W$ of system parameters: $\widehat{N} = J/S = W(v_0, x_0, \ldots)$, where $S$ denotes S when $\mathrm{d}S/\mathrm{d}t = 0$ and inputs to $W$ are the two or more parameters of $P_x(N)$ and $P_v(N)$ (denoted as $x_i$ and $v_i$, respectively) and parameters $n_i$, should they exist, of the species abundance distribution. Rearranging $J/S = W(v_i, x_i, \ldots)$, it follows that the S-nullclines are linear with a slope determined by $W$:

$$S = J/W(v_i, \varepsilon_i, \ldots) \quad \text{(S2a)}$$

or

$$J = W(v_i, \varepsilon_i, \ldots)S \quad \text{(S2b)}$$

If $P_x(N)$ and $P_v(N)$ intersect more than once, there is more than one equilibrium $\hat{N}$ and so more than one S-nullcline (Box 1). However, for typical extinction and speciation functions, there are only one or two intersections. Additional intersections of the speciation and extinction curves at large scales and across a variety of biota resulting from more complicated functions for $P_v$ and $P_x$ seem unlikely, requiring an assumption of more idiosyncratic eco-evolutionary associations. Thus, here we only consider the baseline expectation that there are one or two S-nullclines.

Note that the existence of a diversity equilibrium requires that $P_x(N)$ and $P_v(N)$ intersect at a positive abundance, because otherwise communities can never have a positive equilibrium richness since speciation would be greater than extinction across all abundance classes or vice versa. Similarly, $f_N$ must be able to be sufficiently positive both over some range of *N* where $P_x(N) \geq P_v(N)$ and where $P_x(N) < P_v(N)$ so that there exists a mean abundance that allows $X_{TOT} = V_{TOT}$. For instance, if the assumed SAD function has an upper tail cutoff point (that is, $f_N(N > N_{max}) = 0$), at abundance $N_{max}$ less than the intersection of the extinction and origination curves (assuming they only intersect once) then the system could never achieve an equilibrium. It's also worth noting that if $P_v(N)$ and $P_x(N)$ are identical, any mean abundance allows for $\mathrm{d}S/\mathrm{d}t = 0$ and the function *W* does not exist; however, due to the contrasting mechanics of speciation and extinction, this scenario is impossible, since at low *N*, extinction probability is bound to be very high, approaching 1, whereas speciation probability is much lower than 1.

To understand the *S*-nullcline behavior, consider the idealized scenario in which all species have the same abundance—the "same-abundance distribution". Solving equation S1 using same-abundance distribution (i.e., a Dirac delta function), it follows that $\widehat{N}$ occurs at the intersection point $\dot{N}$ of $P_v(N)$ and $P_x(N)$. Departures from the same-abundance SAD push $\widehat{N}$ away from $\dot{N}$, generally to larger values, but $\dot{N}$ is the main determinant of $\widehat{N}$. $P_v(N)$ and $P_x(N)$ affect the S and J-dependence of $X_{TOT}$ and $V_{TOT}$, determining when attractor ("stabilizing") or repulsive ("destabilizing") dynamics around the S-nullcline are expected. Specifically, the slopes at $\dot{N}$ determine whether or not the associated S-nullcline is repulsive or an attractor—that is, whether or not following deviations from $\widehat{N}$, changes in S eventually return $N$ to $\widehat{N}$ (*attractor S-nullcline*) or push $N$ further from $\widehat{N}$ (*repulsive S-nullcline*). For the same-abundance SAD, if $dP_x(\dot{N})/dN < dP_v(\dot{N})/dN$, then the S-isocline exhibits attractor dynamics; whereas if $dP_x(\dot{N})/dN > dP_v(\dot{N})/dN$, then repulsive dynamics are exhibited.

*Notes on the Species Abundance Distribution*

To address the effects of the species abundance distribution, we have treated species abundance *N* as a continuous variable, as unless community abundance *J* is exceedingly small, use of a probability density function to characterize the species abundance distribution can work well as a continuous approximation of a discrete variable.

We assume that following any perturbation to an equilibrium state, the species abundance distribution recovers to the original function $f_N(N)$ by the end of the time interval, although its mean may change if the community's diversity or abundance is out of equilibrium. In other words, we assume that the ecological and statistical mechanisms of community assembly and population dynamics act to re-equilibrate the SAD (Pueyo et al. 2007). This is a reasonable baseline assumption, given that if it were broken, the SAD could diverge to an extreme distribution unless some other mechanisms were invoked to stabilize it into some other form. In several existing ecological theories this continual re-equilibration of the SAD is implicitly assumed or it is the result of modeled population dynamics, such as in the neutral theory of biodiversity (Hubbell 2001). However, we have not assumed any constraints on the population dynamics of individual species, only that the results of the ensemble of dynamics follows our functions for extinction, speciation, and the SAD. We do not assume that communities far from equilibrium have SADs identical to $f_N(N)$; however, if their SADs differ, we assume that biodiversity dynamics driven by differences between extinction and origination are the dominant drivers of d$S$/d$t$. This assumption is warranted, at least for making coarse-grained predictions of equilibrium stability and diversity trajectories, as we show in Figure 4 that the SAD has a minimal effect on biodiversity patterns (compared to other variables).

*On general features of $P_v(N)$ and $P_x(N)$*

It is worth noting there are some fundamental biological constraints to $P_x(N)$ and $P_v(N)$ that could be important when modeling particular scenarios involving hyper-rich communities or communities with very small abundances. For asexually reproducing taxa, $P_v(N < 1) = 0$ and $P_x(N < 1) = 1$, whereas for sexually reproducing taxa, $P_v(N < 2) = 0$ and $P_x(N < 2) = 1$ (since a couple is required for sexual reproduction). However, since generally $\widehat{N} \gg 2$ (e.g., in the tree data, it's $\widehat{N} \approx 10^8$ individuals in $10^5$ km$^2$), these constraints are of limited importance for general biodiversity patterns.

Some coarse-grained features of the extinction and speciation curves can be deduced in order to shed light on the determinants of the intersection of curves. Extinction probability generally decreases with *N* due to the reduced effects of demographic and environmental stochasticity on large populations and the detrimental effects of genetic drift in small populations (Rosenzweig 1995; Ovaskainen and Meerson 2010). Extinction probability may, however, stop decreasing or even increase at very large population sizes due to large-scale perturbations that wipe out entire populations, regardless of their size. For example, large populations could be susceptible to epidemics due to infectious diseases being positively population-size dependent.

In contrast, speciation probability may increase or decrease with population size, depending on the geographic template and characteristics of the biota (e.g., traits increasing the chances of allopatric or sympatric speciation). At very low *N*, a population necessarily has a low probability of speciation for both genetic and geographic reasons (Rosenzweig 2001; Gavrilets 2003), whereas the population necessarily has a high probability of extinction. However, unless the biota is on an evolutionary path to full extinction, the speciation curve must be higher than the extinction curve at some larger values of *N*, thereby intersecting the extinction curve and having a higher slope (less negative or positive) than the extinction curve at their intersection. Past this intersection point, with increasing *N*, the speciation curve may: (1) continue to increase with *N*, due to covarying organism traits and geographic factors contributing to speciation likelihood or the per capita probability of speciation being invariant of population size, as in the neutral theory of biodiversity, which implies $P_v(N) \sim N^1$ (Allen and Savage 2007); (2) be relatively invariant of *N*; or (3) eventually decrease with *N* and possibly intersect $P_x(N)$ again–e.g., if more abundant, wide-ranging species tend to be generalists and better dispersers and so less sensitive to geographic barriers (Gaston and Chown 1999; Jablonski and Roy 2003).

Additional intersections of the curves, although theoretically possible, are unlikely, especially at large scales and across a variety of biomes and biota.

*Derivations of* $P_v(N)$ *and* $P_x(N)$

We derive a baseline function for $P_x(N)$ by assuming two general results of stochastic extinction theory that are predicted by a variety of models (Ovaskainen and Meerson 2010): (1) population/species extinction times $t_x$ follow an exponential distribution with the probability density function $f_x(t_x) = (1/T)e^{-(1/T)t_x}$, where $T$ is the mean time to extinction; (2) when the effects of environmental stochasticity dominates over demographic stochasticity (as expected over biogeographic scales), environmental stochasticity generally leads to *T* exhibiting power law behavior when the population size carrying capacity K >>1, such that $T \sim K^{x_1}$, where $x_1$ is the scaling exponent (Ovaskainen and Meerson 2010). Since generally *N* ~ *K* for *K* >> *1* (Ovaskainen and Meerson 2010), we can assume $T = {x_0}^{-1} N^{x_1}$, where $x_0$ is a prefactor, with larger values leading to decreased times to extinction. Consequently, the proportion of species of abundance *N* that go extinct within the time interval Δt is determined by substituting $T = {x_0}^{-1} N^{x_1}$ into $f_x(t_x) = (1/T)e^{-(1/T)t_x}$ and integrating over Δt (i.e., $P_x(N) = \int_0^{\Delta t} x_0 N^{-x_1} e^{-x_0 N^{-x_1} t_x} dt_x$), giving

$$P_x(N) = 1 - e^{-x_0 \Delta t N^{-x_1}} \tag{S3}$$

Since we have integrated over the probability density function $f_x(t_x)$, $P_x(N)$ may be interpreted as the probability that a species of abundance *N* goes extinct in the time interval. Although equation S3 is expected to only rigorously apply to *N* >> 1, prediction error at low *N* should be marginal, given that all stochastic extinction models predict high probabilities of extinction at low *N*.

For $x_1 > 0$, as $N \to \infty$ or $\Delta t \to 0$, equation S3 follows exactly a power law with scaling exponent $x_1$, which can be derived by substituting in $e^y = N$ and solving for $d(log\ P_x\ )/dy$ in the limits of $N \to \infty$ or $\Delta t \to 0$. The prefactor $b$ for this power law ($bN^{-x_1}$) can similarly be derived by using l'Hopital's Rule to solve for $b$ in the limit $N \to \infty$ at the intersection of equation S3 and $bN^{-x_1}$, i.e., $b = \lim_{N\to\infty}(1 - e^{-x_0 N^{-x_1}\Delta t})N^{x_1}$ , which gives $b = \Delta t x_0$. Note that at very small $N$ (the exact values of which depends on $\Delta t$), the power function overestimates $P_x(N)$ compared to equation S3 and can become improper (predicting $P_x$>1 at $N < N_{crit}$). However, since $\Delta t$ can be made arbitrarily small in our theory, the power law can provide a very accurate approximation for $N > N_{crit}$, and for $N < N_{crit}$, $P_x(N) \approx 1$ with reasonably low error. So, in sum, Eq S3 is very well approximated by a power law, which it converges exactly to for $N \to \infty$ or $x_0\Delta t \to 0$, giving

$$P_x(N) \approx \Delta t x_0 N^{-x_1}. \tag{S4}$$

Use of such a power law can help facilitate development of analytical predictions and heuristic understanding. We verified that using equation S4 provides nearly identical results regarding the relationships between variables (such as *S* and *J*) to equation S3 for a range of ecologically and evolutionary relevant parameter values (Figure 3 and Figure S1).

Developing baseline theory for speciation rate's dependence on *N* is less straightforward, as there's much more limited theoretical understanding of it. An integrative consideration of population genetics, biogeography, and evolutionary ecology suggests that population size and its correlates could have both positive and negative effects on rates of speciation. For instance, large populations tend to have large geographic ranges, which could increase the chance that a population is separated by a geographic barrier that leads to allopatric speciation (Rosenzweig 1995; Smyčka et al. 2023). Similarly, higher *N* could increase the chance that an individual is born with a mutation that leads to sympatric speciation or a trait that allows the individual to occupy a new niche or area. On the other hand, species with small *N* may tend to have certain traits (such as being ecological specialists), that may make them more likely to speciate. For example, if they are obligate mutualists (like pollinators feeding on nectar) or parasites, then there may be forces to coevolve with their hosts—the speciation of the host eventually being accompanied by the speciation of the pollinator or parasites as the populations coevolve with the new host species (reflecting diversity begetting diversity).

Given the uncertainty, complexity, and idiosyncrasy surrounding the speciation process, we adopt a scaling approach identical to our approach to extinction, such that $T = {v_0}^{-1} N^{-v_1}$, where $v_0$ is a prefactor, with larger values leading to decreased mean times to speciation (and thus increased mean probabilities of speciation). This gives $P_v(N) = 1 - e^{-v_0 \Delta t N^{v_1}}$ and thus the scaling approximation

$$P_v(N) \approx \Delta t v_0 N^{v_1} \quad \text{(S5)}$$

Equation S5 can capture the scenario in which the balance of positive and negative effects of *N* on speciation lead to speciation rate increasing or decreasing with *N*. It can be used to address idealized expectations for speciation, including the scenario in which speciation is independent of *N* (i.e., $v_1 = 0$) and the scenario in which speciation rate increases linearly with *N* (i.e., $v_1 = 1$), due to speciation being considered a neutral per capita process, as in the neutral theory of biodiversity (Allen and Savage 2007).

*References*

Allen, A. P., and V. M. Savage. 2007. Setting the absolute tempo of biodiversity dynamics. Ecology letters 10:637–646.

Gaston, K. J., and S. L. Chown. 1999. Geographic range size and speciation. Evolution of biological diversity 236–259.

Gavrilets, S. 2003. Perspective: models of speciation: what have we learned in 40 years? Evolution 57:2197–2215.

Hubbell, S. P. 2001. The unified neutral theory of biodiversity and biogeography Princeton University Press. Princeton, New Jersey, USA.

Jablonski, D., and K. Roy. 2003. Geographical range and speciation in fossil and living molluscs. Proceedings of the Royal Society of London. Series B: Biological Sciences 270:401–406.

Ovaskainen, O., and B. Meerson. 2010. Stochastic models of population extinction. Trends in ecology & evolution 25:643–652.

Rosenzweig, M. L. 1995. Species diversity in space and time. Cambridge Univ Pr.

Rosenzweig, M. L. 2001. Loss of speciation rate will impoverish future diversity. Proceedings of the National Academy of Sciences 98:5404–5410.

Smyčka, J., A. Toszogyova, and D. Storch. 2023. The relationship between geographic range size and rates of species diversification. Nature Communications 14:5559.

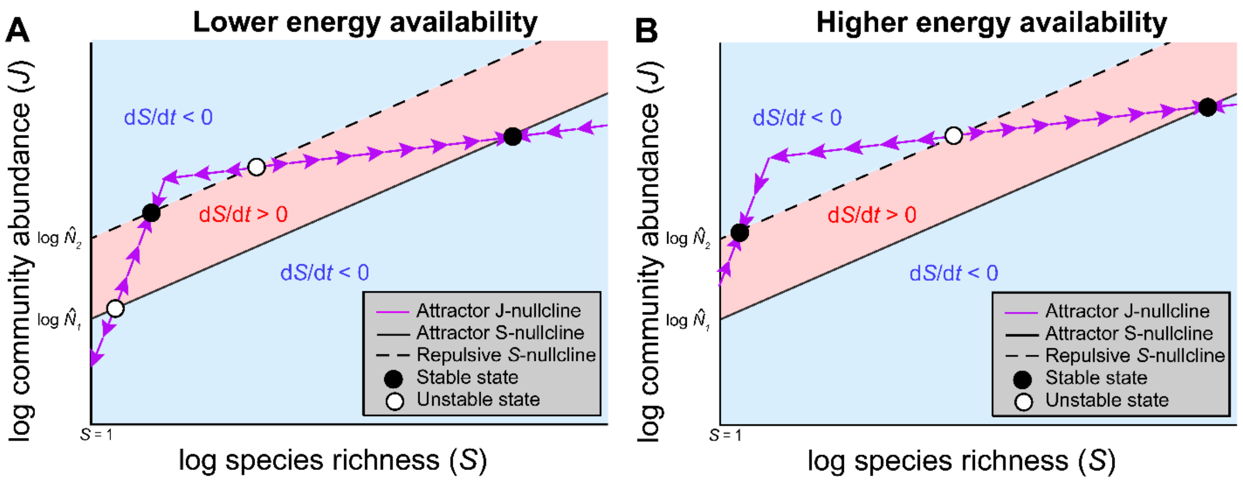


**Figure S1.** Illustration of how an increase in the height of the J-nullcline differentially influences the species richness and community abundance of equilibrium points, depending on the scaling of the J-nullcline around the equilibrium point. Increased energy availability *(E)*, reduced individual metabolic rate *(B)*, or an increase in the ability of communities of a given *S* to use available resources (*c*) increases the height of the J-nullcline, as shown in **B**. When the J-nullcline is steeper than the S-nullcline around the equilibrium point (i.e., if $\beta > 1$ due to facilitation), the unstable and stable equilibrium point's *S* and *J* decreases, as shown in **B.** If the J-nullcline height is sufficiently increased, these equilibrium points may even disappear, due to the facilitation-phase of the J-nullcline no longer intersecting the S-nullclines. In contrast, an increase in the height of the J-nullcline leads to an increase in the *S* and *J* of the stable equilibrium point found at the intersection with the attractor S-nullcline, as well as an increase in the *S* and *J* of the unstable equilibrium point found at the intersection with the repulsive S-nullcline.

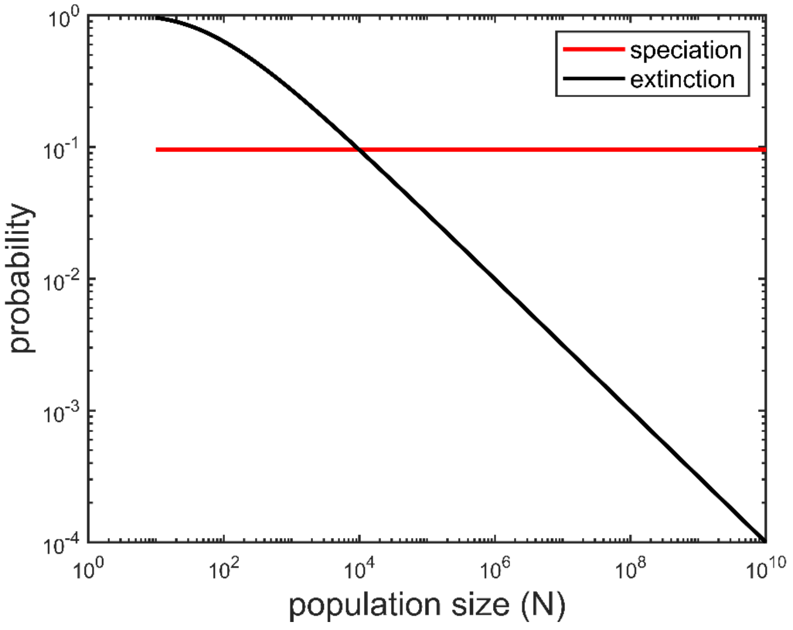


**Figure S1.** Illustration of the population size-dependence of probabilities of extinction and speciation with the exponent parameters assumed for the full model shown in the first row of Figure 4. Because the curves are straight at and near their intersection, the power law approximation works well to model gradients in equilibrium diversity and abundance. Here the speciation probability was assumed to be independent of $N$ (that is, $v_1 = 0$). The plotted speciation curve extends to $N = 1$, but it becomes numerically more intensive to plot with the full model, so was not shown here.

**Table S1.** Results of the Reduced Major Axis regression analysis using a biome-realm combinations' mean country-level species richness as a measure of regional species richness for that biome-realm combination (i.e., without standardizing to 100,000 $km^2$).

| Y | X | Observed Slope | Lower 95% CI | Upper 95% CI | Intercept | $R^2$ |
|---|---|---|---|---|---|---|
| ln regional S | absolute latitude | -0.084 | -0.109 | -0.060 | 8.525 | 0.43 |
| ln local S (per 0.1 $km^2$) | ln regional S | 1.111 | 0.615 | 1.607 | -2.984 | 0.75 |
| ln local S (per 0.01 $km^2$) | ln regional S | 1.08 | 0.707 | 1.454 | -3.518 | 0.79 |

**Table S2**. Results of the Reduced Major Axis regression analysis of inverse mean annual air temperature (1/kT, where k is Boltzmann's constant in eV and T is temperature in kelvins), log species richness (*S*), log total number of individuals (*J*), log mean population abundance ($\widehat{N}$) in trees from terrestrial biomes.

| Y | X | Observed Slope ("Activation Energy") | Lower 95% CI | Upper 95% CI | Intercept | $R^2$ | Area ($km^2$) |
|---|---|---|---|---|---|---|---|
| ln $S$ | $1/kT$ | -1.240 | -1.777 | -0.703 | 54.350 | 0.765 | 0.1 |
| ln $S$ | $1/kT$ | -1.041 | -1.377 | -0.704 | 44.765 | 0.843 | 0.01 |
| ln $\widehat{N}$ | $1/kT$ | 0.981 | 0.576 | 1.385 | -34.361 | 0.787 | 0.1 |
| ln $\widehat{N}$ | $1/kT$ | 0.886 | 0.569 | 1.203 | -32.562 | 0.807 | 0.01 |
| ln $J$ | $1/kT$ | -0.333 | -0.562 | -0.105 | 22.986 | 0.385 | 0.1 |
| ln $J$ | $1/kT$ | -0.257 | -0.421 | -0.093 | 16.226 | 0.385 | 0.01 |

**Table S3.** Calculated/estimated mean tree species richness and community abundance of different biomes and biogeographic realms at spatial grains ranging from 0.01 $km^2$ to 100,000 $km^2$.

| Scale ($km^2$) | Habitat-realm combination | Counts | Mean $log_{10}$ species richness | Mean $log_{10}$ community abundance | Mean annual temperature (ºC) | Mean latitude (º) |
|---|---|---|---|---|---|---|
| 100000 | Boreal Forests/Taiga_Nearctic | 1 | 1.375 | NA | -7.059 | 57.715 |
| 100000 | Boreal Forests/Taiga_Palearctic | 4 | 1.375 | NA | -0.406 | 63.304 |
| 100000 | Deserts and Xeric Shrublands_Afrotropic | 4 | 2.224 | NA | 21.478 | 21.268 |
| 100000 | Deserts and Xeric Shrublands_Australasia | 1 | 2.787 | NA | 21.458 | 25.574 |
| 100000 | Deserts and Xeric Shrublands_Nearctic | 5 | 2.201 | NA | 11.375 | 36.906 |
| 100000 | Deserts and Xeric Shrublands_Neotropic | 7 | 2.714 | NA | 24.488 | 8.424 |
| 100000 | Deserts and Xeric Shrublands_Palearctic | 20 | 1.462 | NA | 18.844 | 31.057 |
| 100000 | Flooded Grasslands and Savannas_Afrotropic | 1 | 2.541 | NA | 21.223 | 13.453 |
| 100000 | Mediterranean Forests, Woodlands and Scrub_Neotropic | 1 | 1.797 | NA | 8.646 | 35.513 |
| 100000 | Mediterranean Forests, Woodlands and Scrub_Palearctic | 9 | 1.844 | NA | 15.131 | 37.344 |
| 100000 | Montane Grasslands and Shrublands_Afrotropic | 3 | 2.212 | NA | 19.175 | 23.833 |
| 100000 | Montane Grasslands and Shrublands_Palearctic | 4 | 2.705 | NA | 2.198 | 35.633 |
| 100000 | Temperate Broadleaf and Mixed Forests_Nearctic | 24 | 2.297 | NA | 10.757 | 39.567 |
| 100000 | Temperate Broadleaf and Mixed Forests_Palearctic | 38 | 2.085 | NA | 8.902 | 44.296 |
| 100000 | Temperate Conifer Forests_Indo-Malay | 1 | 2.772 | NA | 10.254 | 27.416 |
| 100000 | Temperate Conifer Forests_Nearctic | 9 | 2.132 | NA | 11.992 | 37.441 |
| 100000 | Temperate Conifer Forests_Palearctic | 5 | 2.111 | NA | 7.122 | 43.913 |
| 100000 | Temperate Grasslands, Savannas and Shrublands_Nearctic | 9 | 1.994 | NA | 10.198 | 40.904 |
| 100000 | Temperate Grasslands, Savannas and Shrublands_Neotropic | 1 | 2.275 | NA | 14.200 | 35.178 |
| 100000 | Temperate Grasslands, Savannas and Shrublands_Palearctic | 4 | 1.511 | NA | 5.282 | 40.460 |
| 100000 | Tropical and Subtropical Coniferous Forests_Indomalay | 1 | 2.621 | NA | 14.053 | 28.253 |
| 100000 | Tropical and Subtropical Coniferous Forests_Neotropic | 2 | 3.060 | NA | 24.002 | 14.277 |
| 100000 | Tropical and Subtropical Dry Broadleaf Forests_Indomalay | 1 | 3.059 | NA | 26.121 | 15.145 |
| 100000 | Tropical and Subtropical Dry Broadleaf Forests_Neotropic | 1 | 3.368 | NA | 25.008 | 8.514 |
| 100000 | Tropical and Subtropical Grasslands, Savannas and Shrublands_Afrotropic | 27 | 2.493 | NA | 24.901 | 10.079 |
| 100000 | Tropical and Subtropical Grasslands, Savannas and Shrublands_Neotropic | 12 | 2.823 | NA | 23.602 | 14.275 |
| 100000 | Tropical and Subtropical Moist Broadleaf Forests_Afrotropic | 6 | 2.850 | NA | 24.606 | 3.482 |
| 100000 | Tropical and Subtropical Moist Broadleaf Forests_Indomalay | 12 | 3.035 | NA | 21.312 | 22.793 |
| 100000 | Tropical and Subtropical Moist Broadleaf Forests_Neotropic | 22 | 3.006 | NA | 23.277 | 12.519 |
| 100000 | Tropical and Subtropical Moist Broadleaf Forests_Palearctic | 2 | 3.411 | NA | 15.457 | 25.896 |
| 0.1 | Boreal Forests/Taiga_Nearctic | 1 | 1.176 | 3.884 | 4.200 | 45.290 |
| 0.1 | Temperate Broadleaf and Mixed Forests_Nearctic | 4 | 1.586 | 4.098 | 10.000 | 40.640 |
| 0.1 | Temperate Broadleaf and Mixed Forests_Palearctic | 5 | 1.409 | 3.979 | 7.640 | 42.502 |
| 0.1 | Temperate Conifer Forests_Nearctic | 2 | 0.972 | 4.026 | 8.450 | 41.790 |
| 0.1 | Tropical and Subtropical Dry Broadleaf Forests_Indomalay | 1 | 2.310 | 4.195 | 23.600 | 14.430 |
| 0.1 | Tropical and Subtropical Moist Broadleaf Forests_Afrotropic | 2 | 2.466 | 4.296 | 24.700 | 3.255 |

| | | | | | | |
|---|---|---|---|---|---|---|
| 0.1 | Tropical and Subtropical Moist Broadleaf Forests_Indomalay | 10 | 2.268 | 4.195 | 22.010 | 16.473 |
| 0.1 | Tropical and Subtropical Moist Broadleaf Forests_Neotropic | 3 | 2.596 | 4.207 | 22.667 | 2.373 |
| 0.1 | Tropical and Subtropical Moist Broadleaf Forests_Palearctic | 1 | 2.270 | 4.411 | 11.100 | 29.770 |
| 0.01 | Deserts and Xeric Shrublands_Neotropic | 1 | 1.568 | 2.489 | 23.600 | 10.530 |
| 0.01 | Mangroves_Neotropic | 1 | 2.170 | 2.698 | 26.200 | 1.180 |
| 0.01 | Temperate Broadleaf and Mixed Forests_Palearctic | 3 | 0.911 | 2.544 | 7.900 | 48.639 |
| 0.01 | Temperate Conifer Forests_Palearctic | 14 | 0.667 | 2.498 | 5.500 | 48.990 |
| 0.01 | Tropical and Subtropical Dry Broadleaf Forests_Neotropic | 14 | 1.906 | 2.781 | 23.700 | 13.629 |
| 0.01 | Tropical and Subtropical Grasslands, Savannas and Shrublands_Afrotropic | 31 | 1.660 | 2.581 | 24.377 | 3.720 |
| 0.01 | Tropical and Subtropical Grasslands, Savannas and Shrublands_Neotropic | 22 | 1.575 | 2.772 | 26.109 | 12.720 |
| 0.01 | Tropical and Subtropical Moist Broadleaf Forests_Afrotropic | 103 | 1.888 | 2.619 | 24.730 | 3.404 |
| 0.01 | Tropical and Subtropical Moist Broadleaf Forests_Indomalay | 107 | 1.691 | 2.726 | 24.537 | 14.101 |
| 0.01 | Tropical and Subtropical Moist Broadleaf Forests_Neotropic | 177 | 2.119 | 2.802 | 25.759 | 6.321 |
